# Integrating large-scale neuroimaging research datasets: harmonisation of white matter hyperintensity measurements across Whitehall and UK Biobank datasets

**DOI:** 10.1101/2020.07.28.208579

**Authors:** Valentina Bordin, Ilaria Bertani, Irene Mattioli, Vaanathi Sundaresan, Paul McCarthy, Sana Suri, Enikő Zsoldos, Nicola Filippini, Abda Mahmood, Luca Melazzini, Maria Marcella Laganà, Giovanna Zamboni, Archana Singh-Manoux, Mika Kivimäki, Klaus P Ebmeier, Giuseppe Baselli, Mark Jenkinson, Clare E Mackay, Eugene P Duff, Ludovica Griffanti

**Author notes:** These authors contributed equally to this work (Eugene P Duff, Ludovica Griffanti). Corresponding author: Ludovica Griffanti, Wellcome Centre for Integrative Neuroimaging (WIN), Department of Psychiatry, Warneford Ln, Headington, Oxford, OX3 7JX.

## Abstract

Large scale neuroimaging datasets present the possibility of providing normative distributions for a wide variety of neuroimaging markers, which would vastly improve the clinical utility of these measures. However, a major challenge is our current poor ability to integrate measures across different large-scale datasets, due to inconsistencies in imaging and non-imaging measures across the different protocols and populations. Here we explore the harmonisation of white matter hyperintensity (WMH) measures across two major studies of healthy elderly populations, the Whitehall II imaging sub-study and the UK Biobank. We identify pre-processing strategies that maximise the consistency across datasets and utilise multivariate regression to characterise sample differences contributing to study-level differences in WMH variations. We also present a parser to harmonise WMH-relevant non-imaging variables across the two datasets. We show that we can provide highly calibrated WMH measures from these datasets with: (1) the inclusion of a number of specific standardised processing steps; and (2) appropriate modelling of sample differences through the alignment of demographic, cognitive and physiological variables. These results open up a wide range of applications for the study of WMHs and other neuroimaging markers across extensive databases of clinical data.

**HIGHLIGHTS:**

- We harmonised measures of WMHs across two studies on healthy ageing
- Specific pre-processing strategies can increase comparability across studies
- Modelling of biological differences is crucial to provide calibrated measures

## INTRODUCTION

The increasing availability of brain MRI datasets through multi-centre studies, consortia, and data sharing platforms, along with the increased power of computational resources, allows for the possibility of merging datasets and achieving unprecedent statistical power (Smith and Nichols, 2018). This has massively increased the range of research questions that can now be tackled. Moreover, this provides the possibility of generating normative distributions of neuroimaging markers, which would vastly improve the clinical utility of these measures. However, the increasing use of combined datasets has raised the important issue of ensuring that measures are consistent across datasets. The process of *harmonisation* aims to remove non-biological variability related to the measurement process, while preserving the biological and especially the clinically-relevant variability present in the data.

In this work we aimed to combine different harmonisation approaches to develop a harmonisation pipeline for MRI-derived measures of white matter hyperintensities (WMHs) of presumed vascular origin (Wardlaw et al., 2013) on two large datasets related to healthy ageing that are part of the Dementias Platform UK (Bauermeister et al., 2020): the Whitehall II imaging sub-study (WH) (Filippini et al., 2014) and the UK Biobank (UKB) (Miller et al., 2016). The first underwent a scanner upgrade during data collection, and therefore contains data acquired on two scanners using the same protocol (Zsoldos et al., 2020). The second is instead a dataset that despite being also focused on the aging population used a different scanner, protocol, and set of non-imaging variables (demographic, cognitive and physiological) from WH. Our goal was to find the best combination of approaches able to reduce differences in WMH measures extracted from these datasets. This would help providing a comprehensive protocol to successfully reduce biases and promote data integration.

The importance of characterising ageing-associated vascular damage is increasingly recognised, since vascular disease contributes to more than half of dementia cases, often in conjunction with Alzheimer’s disease pathology (Arvanitakis et al., 2016; Debette et al., 2010). Among the signs of cerebral small vessel disease (SVD), WMHs are one of the most commonly evaluated, but their underlying pathology and clinical impact on cognition is still poorly understood (Wardlaw et al., 2013), and possibly affected by age (Zamboni et al., 2019). Being able to combine datasets would give further insight on the relationships between WMHs, its risk factors and clinical outcomes. This would not only improve statistical power, but also enable merging complementary information from datasets such as WH and UKB. For example, WH includes detailed longitudinal cognitive and behavioural assessments, while the UKB dataset has a bigger sample size and is more generalisable to the population (wider age range and more even gender balance than WH). An ability to integrate WMH data across these two datasets would open up the ability to gain novel insights into the prognostic value of WMHs.

While many harmonisation approaches have been developed for T1-weighted (e.g. Fortin et al., 2018; Zanadifar et al., 2018) and diffusion MRI (e.g. Fortin et al., 2017; Mirzaalian et al., 2016), harmonisation approaches for T2-weighted scans and the quantification of WMHs (and other lesions) are still lacking, despite the recognition that biases are also present in this modality (Shinohara et al., 2017; Guo et al., 2019). Consortia and working groups (Wardlaw et al., 2013; Smith et al., 2019) recognised the need to standardise the assessment of SVD and proposed a set of standard definitions, acquisition protocols and a framework for developing neuroimaging biomarkers of cerebral small vessel disease. The HARmoNising Brain Imaging MEthodS for VaScular Contributions to Neurodegeneration (HARNESS) initiative (https://harness-neuroimaging.org) also set up web-based repositories of protocols, software tools and rating scales to facilitate multi-centre research. While all these resources contribute to more standardised assessment of WMHs, what is still currently lacking is a way to make quantitative measures truly consistent.

The datasets we selected for this study allow us to test *retrospective* (i.e. after data collection) harmonisation strategies in the presence (WH scanner upgrade) and absence (WH-UKB) of *prospective* (i.e. prior to data collection) harmonisation. Harmonised acquisition protocols are commonly done in consortia and multi-centre studies (Jack et al., 2008; Potvin et al., 2019), as an agreement on a main set of collection procedures and common measures prior to data collection, to facilitate future integration or comparison of data. However, when dealing with MRI-derived measures some retrospective harmonisation is still needed. This is because, even after careful protocol harmonisation, systematic differences in the images can remain (related to scanner vendor, model, non-linearity of imaging gradients, magnetic field homogeneity, signal-to-noise ratio etc.) leading to bias in the MRI-derived measures (Kruggel et al., 2010; Potvin et al., 2019; Shinohara et al., 2017; Mirzaalian et al., 2016; Guo et al., 2019). At the image pre-processing level, harmonisation strategies aim to remove the non-biological variability, leading to more similar images (Mirzaalian et al., 2016; Dewey et al., 2018), and to design/use analysis tools or pipelines that give consistent performance on different datasets and produce well-matched measures (Erus et al., 2018; Zandifar et al., 2018; Guo et al., 2019). At the analysis level, harmonisation approaches aim to make measures consistent across datasets, some also modelling biological variability, so that characteristics of imaging site and study are removed from the data (Fortin et al., 2017; Fortin et al., 2018; Pomponio 2020). Despite rapid progress, MRI data harmonisation remains a challenge. Due to the different nature of the biases involved, a single strategy is unlikely to achieve successfully harmonised data (Wachinger et al., 2019; Glocker et al., 2019).

A key element of our project involved increasing the robustness of FSL-BIANCA, a supervised classification method for segmenting WMHs (Griffanti et al., 2016). Briefly, BIANCA classifies the image voxels based on their intensity and spatial features using the k-nearest neighbour (k-NN) algorithm. The intensity features can be extracted from multiple MRI modalities, making it a very versatile tool. Being a supervised method, it needs some examples of manually segmented WMHs for training the algorithm. The output image represents the probability per voxel of being a WMH and can then be thresholded to obtain the final binary mask (see Griffanti et al., 2016 for further details). BIANCA has been tested on vascular, neurodegenerative and healthy populations. It achieved excellent performance scores with respect to manual annotation and visual ratings and it is also registered among the software tools on the HARNESS initiative website (https://software.harness-neuroimaging.org/harness-software-catalog/bianca.html).

Starting from the WH ‘scanner upgrade scenario’, we explored the effect of processing choices that are particularly relevant for harmonisation: the impact of the rater performing the manual segmentations used to train BIANCA (likely to be different across datasets), the effect of bias field correction (since the distribution of radio frequency (RF) inhomogeneities is unique to the scanner), the choice of intensity features used to classify WMHs (which also depends on the available modalities for a specific dataset) and the composition of the training set for BIANCA (i.e. from which dataset(s) the example segmentations should be taken). We then extended the evaluation to the ‘retrospective data merging’ scenario, to harmonise WH and UKB. In this case the pipeline also included a non-imaging variables harmonisation step, to be able to model the biological variability in WMHs. Finally, based on the optimal settings obtained, we propose a set of recommendations for improving WMH comparability across datasets.

## METHODS

### Datasets

The datasets we used in this work are WH and UKB.

The first, described in Filippini et al. (2014), is part of a large longitudinal study, namely the Whitehall II Study, that explores the social determinants of health. It involves a sample of British civil servants (age range 60-85 years) who were first recruited in 1985 and participated in a number of phases of clinical/cognitive assessment. Seven hundred and seventy-four participants were selected randomly to receive multi-modal brain MRI scans and a detailed cognitive battery at the Oxford Centre for Functional MRI of the Brain (FMRIB) as part of the Imaging Sub-study (2012-2016). Out of those, we excluded 18 participants with evident brain abnormalities other than WMHs (e.g. tumour, stroke, multiple sclerosis), and 17 due to poor quality of the available MRI scans or lack of some of the MRI contrast of interest. As a result, the analysis was performed on a total number of 739 subjects, of which 528 (WH1) were imaged with a 3T Siemens Verio scanner (SC1) and 211 (WH2) with a 3T Siemens Prisma (SC2). Alongside the WH cohort, 5 additional young and healthy participants (age 31 ± 4.9 years, age range 26-39 years, 2 males) were also scanned at FMRIB on both SC1 and SC2 (‘traveling heads’). Even though these subjects did not have any WMHs, the MRI data allowed us to get additional insight on non-biological sources of variability in the images and test some harmonisation approaches.

The second dataset is the UKB imaging study, a sub-study of the UKB prospective epidemiological study gathering extensive questionnaires, physical and cognitive measures, and biological samples from predominantly healthy participants. The project imaging component (Miller et al., 2016), currently ongoing, aims at collecting detailed diagnostic MRI scans from 100,000 UKB participants. The sample available at the time of our work included 14,503 subjects with scans released by January 2019 (age range 50-80 years). Out of those with available MRI data, we selected 3,205 subjects that had no missing data in the non-imaging variables of interest (details below). This allowed us to avoid performing data imputation, which could have introduced an additional source of variability, while still having a large number of subjects to focus on the methodological goal of imaging data harmonisation. Ten further participants were excluded due to other brain abnormalities. The resulting UKB dataset was therefore composed of 2̇,295 participants.

#### Non-imaging variables

In order to model the biological variability in WMH measures across datasets, we selected non-imaging variables with a potential link to WMHs. An example of such variables is age. It is one of the most important risk factor for WMHs, and WH and UKB have only partially overlapping age ranges (WH: 60-85 years; UKB: 50-80 years). Therefore, we considered particularly important to take it into account as source of biological variability when comparing measures of WMHs across datasets. A total of 33 variables, including demographic, clinical and cognitive factors were selected among those available for the WH dataset. Subsequently, when performing harmonisation between the WH and UKB datasets, we excluded 4 variables due to lack of availability for all participants within the UKB cohort, or due to substantial differences in the data collection across the two datasets (e.g. the design or administration of certain cognitive tests). The full list of non-imaging variables selected for both datasets is presented in Table 1.

**Table 1.**
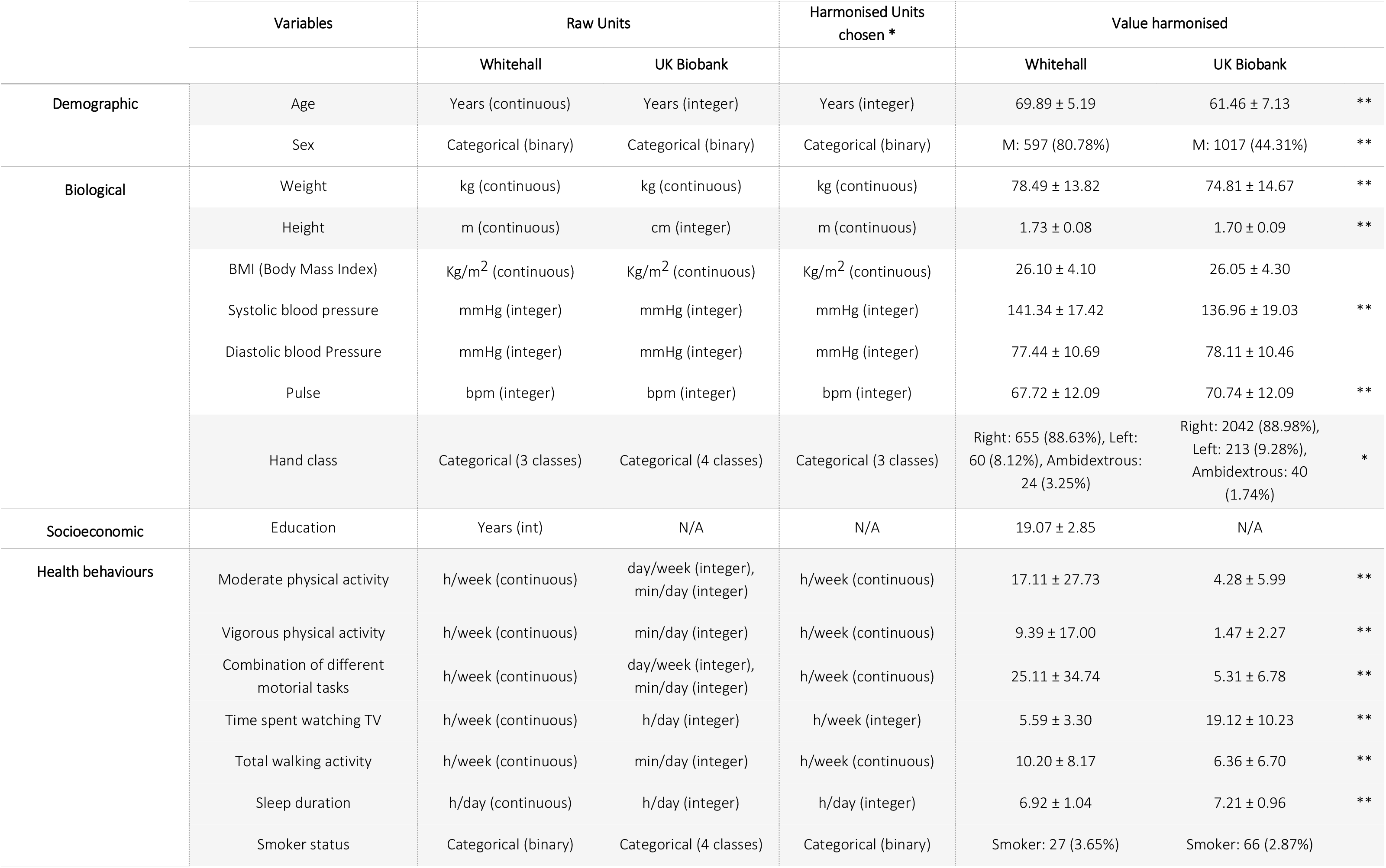
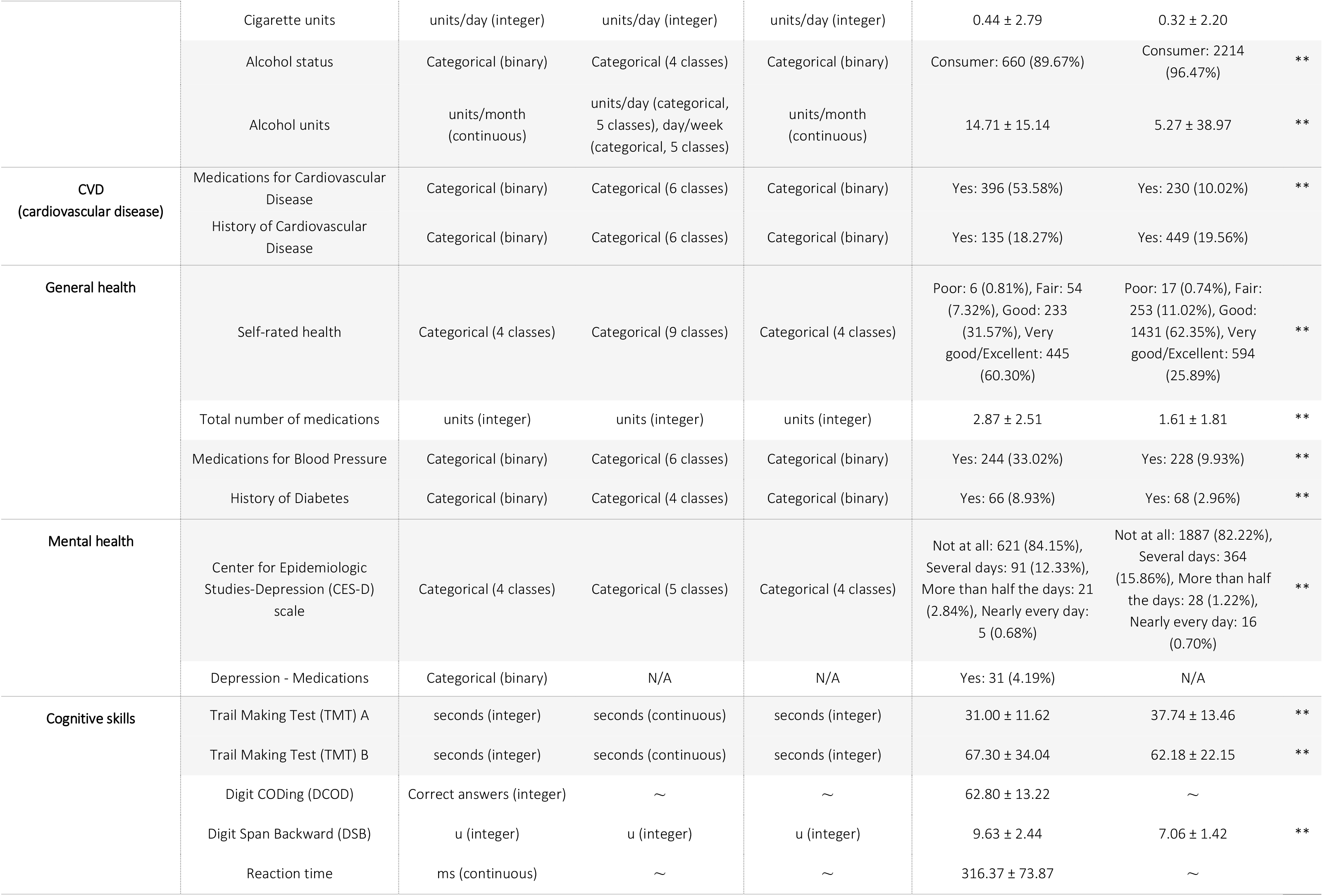
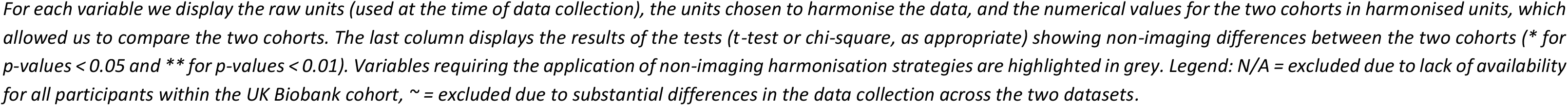
Details of the non-imaging variables selected for our study.

#### MRI data acquisition

Acquisition details for the datasets involved in our analysis are listed in Table 2.

**Table 2.** Acquisition details for the three scanners involved in our study.

|  | Whitehall |  |  |  |  |  | UK Biobank |  |
| --- | --- | --- | --- | --- | --- | --- | --- | --- |
|  | 3T Siemens Verio (WH1) |  |  | 3T Siemens Prisma (WH2) |  |  | 3T Siemens Skyra (UKB) |  |
| Sequence | FLAIR | T1 (MEMPR) | dMRI (EPI) | FLAIR | T1 (MPRAGE) | dMRI (EPI) | FLAIR | T1 |
| TR (ms) | 9000 | 2530 | 8900 | 9000 | 1900 | 8900 | 5000 | 2000 |
| TE (ms) | 73 | 1.79/3.65/<br>5.51/7.37 | 91.2 | 73 | 3.97 | 91 | 395.0 | 2.01 |
| Flip angle (degrees) | 150 | 7 | ----- | 150 | 8 | ----- | ----- | 8 |
| Voxel dimension (mm<sup>3</sup>) | 0.9x0.9x3 | 1x1x1 | 2x2x2 | 0.4x0.4x3 | 1x1x1 | 2x2x2 | 1.05x1x1 | 1x1x1 |
| FoV read (mm) | 220 | 256 | 192 | 220 | 192 | 192 | 256 | 256 |
| FoV phase (%) | 100 | 100 | 100 | 100 | 100 | 100 | 100 | 100 |
| Base resolution | 256 | 256 | 96 | 256 | 256 | 96 | 256 | 256 |
| Phase resolution (%) | 100 | 100 | 100 | 100 | 100 | 100 | 100 | 100 |
| TI (ms) | 2500 | 1380 | ----- | 2500 | 904 | ----- | 1800 | 880 |
| Bandwidth (Hz/Px) | 283 | 651 | 1680 | 283 | 200 | 1680 | 781 | 240 |
| Orientation | Transversal | Sagittal | Transversal | Transversal | Transversal | Transversal | Sagittal | Sagittal |
| b-value (s/mm<sup>2</sup>) | ----- | ----- | 1500 | ----- | ----- | 1500 | ----- | ----- |
| Directions (n.) | ----- | ----- | 60 + 6 b0 (1 reversed PE) | ----- | ----- | 60 + 6 b0 (1 reversed PE) | ----- | ----- |
| Acquisition time | 4m 14s | 6m 12s | 9m 56s | 4m 14s | 5m 31s | 10m 41s | 5m 52s | 4m 54s |

Legend: FLAIR, fluid attenuated inversion recovery; MEMPR, Multi-Echo MPRAGE; MPRAGE, Magnetization Prepared Rapid Acquisition Gradient Echo; dMRI, diffusion MRI; EPI, Echo Planar Imaging; TR, repetition time; TE, echo time; FoV, field of view; TI, inversion time; PE, Phase Encoding.

For the WH study, two MRI scanners were used, due to the scanner upgrade two-thirds of the way through the study: a 3T Siemens Magnetom Verio scanner (SC1) with a 32-channel receive head coil (April 2012 – December 2014) and a 3T Siemens Prisma scanner (SC2) with a 64-channel receive head-neck coil in the same centre (July 2015 – December 2016). The MRI modalities used for WMH segmentation were Fluid Attenuated Inversion Recovery (FLAIR) scans, T1-weighed scans and diffusion-weighted scans (dMRI), to derive Fractional Anisotropy (FA) maps. The MRI sequence parameters were either identical or closely matched between the two scanners.

For the UKB dataset, MRI acquisition was carried out using a 3T Siemens Skyra with a 32-channel receive head coil (full details in Miller et al., 2016). As regards the MRI modalities, for the current study we used FLAIR scans and T1-weighed scans. We decided not to include dMRI within the WMH quantification pipeline, because the requirement to have 3 usable MRI modalities for each subject would have caused the exclusion of a small, yet significant amount of data (see (Alfaro-Almagro et al., 2018) for an indication of usable data for each modality). In fact, currently released measures of WMHs for UKB are extracted using T1-weighted and FLAIR only. Moreover, unlike T1-weighted and FLAIR scans, dMRI with 6 or more directions (needed to perform Diffusion Tensor Imaging and generate FA maps) are not very common in clinical contexts. Therefore, being able to obtain consistent WMH estimates with common sequences would make our approach more widely applicable.

#### MRI pre-processing

All the available MRI scans underwent pre-processing using FSL v.6.0 tools (Jenkinson et al., 2012) before being fed to BIANCA for WMH segmentation. T1-weighted scans were processed using fsl_anat, which performs bias correction, brain extraction, and partial-volume tissue segmentation using FAST (Zhang et al., 2001). The sum of the volumes for the three tissue classes was used as total brain volume to normalise WMH measures. We used an exclusion mask for cortical grey matter and structures that can appear hyperintense on FLAIR and for which BIANCA is not currently optimised (putamen, globus pallidus, nucleus accumbens, thalamus, brainstem, cerebellum, hippocampus, amygdala) (details in (Griffanti et al., 2016)). FLAIR images were brain-extracted using BET (Smith et al., 2002) and bias field corrected with FAST (Zhang et al., 2001). Images without bias field correction were also used to evaluate the effect of this pre-processing step on the WMH measures. For WH data, dMRI scans were pre-processed as described in (Filippini et al., 2014) and a diffusion tensor model was fit at each voxel to obtain FA maps.

Since BIANCA works in single-subject space, we used FLIRT (Jenkinson et al., 2001) to register all the MRI modalities to the FLAIR scan, chosen as reference modality. Then, we masked the latter with the exclusion mask derived from the T1-weighted images. The transformation between FLAIR and MNI space for each subject was also calculated (using FLIRT) to be used by BIANCA to derive the spatial features (MNI coordinates).

As BIANCA requires several parameter choices, we tested the influence of those that are particularly relevant for harmonisation, while keeping the others constant. We performed a preliminary analysis to assess the best combination of settings that produced consistent performances for segmentation accuracy and specificity across datasets. The best settings were found to be in line with the suggested parameters in (Griffanti et al., 2016) and previously used in studies using BIANCA on the WH dataset (Griffanti et al., 2018). Therefore, we fixed the following parameters for BIANCA throughout our study: 2,000 training points representing WMH lesions, 10,000 points representing non-lesion voxels, a patch size of dimension 3 voxels and a spatial weighting coefficient equal to 2.

A subset of manually segmented WMH images was available from each dataset to train BIANCA and to evaluate its segmentation performance in a cross-validated manner. The segmented data included 24 participants from the WH1 dataset, 24 from the WH2 dataset and 12 from the UKB dataset. The 24 subjects from WH1 were manually annotated by two raters (R1, R2). Rater 2 repeated their annotation a year later (R2a, R2b) enabling us to assess the effects of within- and between-rater variability on the WMH measures. Rater 2 also labelled the 24 scans from the WH2 dataset. For UKB we used the manual masks of 12 subjects used in the released imaging pipeline (Alfaro-Almagro 2018).

### Harmonisation pipeline

During our work we dealt with two scenarios: the first aimed at harmonising the two Whitehall imaging sub-studies (WH1 and WH2) representing data before and after the scanner upgrade within the same cohort and centre; the second addressed the integration of different cohorts (WH and UKB) acquired on different scanners at different centres. The different scenarios allowed us to test the effect of different factors affecting data and required some changes in the harmonisation pipelines applied.

#### Scanner upgrade (Whitehall)

We started the analysis with the scanner upgrade scenario (WH1 and WH2) that included prospective harmonisation in the study design: the same non-imaging variables were collected and the MRI protocol was as close as possible for the two scanners. Retrospective harmonisation was therefore not needed for the non-imaging data but carried out on the images.

The availability of manual masks from multiple raters, ‘traveling heads’ data and FA maps for most of the participants allowed us to study the impact of: (i) the rater performing the manual labelling, (ii) the process of bias field correction on FLAIR images, (iii) the composition of the dataset used to train BIANCA (training set) and (iv) the inclusion of FA as one of the MRI modalities used by the segmentation tool to derive intensity features. We compared one option at a time using the metrics described in the *Evaluation metrics* section, while keeping the others fixed, in order to understand how each one could influence the results. We then identified optimal pre-processing and analysis strategies to reduce non-biological variability across datasets, while retaining or taking into account (modelling) the biological variability.

- **Effect of rater**: in the training phase, BIANCA requires manually delineated WMH masks, which are known to suffer from inter- and intra-rater variability (Guo et al., 2019). We wanted to assess whether BIANCA trained with different manual masks (either multiple annotations by different raters or repeated annotations by the same rater) generates WMH segmentations that are more or less variable than the manual annotations among themselves. If BIANCA produced more consistent WMH masks than manual operators, the use of this automated segmentation tool would be advisable to obtain more consistent results. We evaluated this on data from a single scanner (WH1). We had multiple annotations for 24 MRI scans (two raters - R1, R2; and two annotations by R2 one year apart – R2a, R2b – corresponding manual masks M1, M2a, M2b). Between-rater (M1 vs M2a; M1 vs M2b) and within-rater (M2a vs M2b) agreement was calculated in terms of overlap between the manual masks using Dice Similarity Index (DI – see Griffanti et al., 2016). Each set of ratings was then used to train BIANCA and the automated WMH masks (B1, B2a, B2b) were generated using a leave-one-out approach. We then calculated between-rater (B1 vs B2a; B1 vs B2b) and within-rater (B2a vs B2b) agreement also on the automatically segmented masks using DI. Finally, we compared DI values using paired t-tests to assess whether consistency within the automatic WMH segmentations was higher or lower with respect to consistency within the manually labelled masks.
- **Effect of bias field correction**: we assessed the impact of bias field correction (BC) in multiple ways. One indication of successful harmonisation is that harmonised images should be more similar to each other. We evaluated this aspect on the ‘traveling heads’ data available for the WH dataset. Corresponding scans from each of the 5 subjects were first registered to each other and then resampled into the space half-way between the two. We then calculated the cost function (correlation ratio) between the registered images as a measure of image similarity that is not influenced by head position (lower cost function indicates more similar images). The same procedure was repeated on the bias field corrected images. The values of the cost function before and after BC were compared with a paired t-test. Secondly, we investigated the effect of BC on BIANCA performance (i.e. overlap with manual WMH masks) as described in the *Evaluation metrics* section. The manual rater was R2 for both datasets and the training set for BIANCA was the same (24 subjects from WH1). We compared the results obtained before and after BC, to test whether the adoption of this pre-processing step could provide more consistent results across datasets. We then evaluated the effect of BC on the relationship between WMHs and age, and in terms of explained variability of the scanner effect in a multivariate regression model (see *Evaluation metrics* for details).
- **Effect of training set composition for BIANCA:** we compared three different options that could be used to train BIANCA when performing WMH segmentation on multiple datasets: single-site training (using the same training set for all datasets, with examples coming only from one site – 24 subjects from WH1 in our case), site-specific training (training BIANCA on each dataset separately) and mixed training (combining examples from WH1 and WH2, 24 subjects each, in a single training set to apply to all datasets). As before, we exploited several analysis approaches to evaluate which option would lead to better harmonised WMH measures. We investigated the effect of each option on: BIANCA performance, the relationship between WMHs and age, and the weight of the scanner variable in the multivariate regression model. All data were bias field corrected before the analysis (see *Evaluation metrics* for details).
- **Effect of FA information**: as previously mentioned, we did not use FA maps derived from dMRI to inform WMH segmentation for the UKB dataset, but FA maps were used in the WH dataset. Aiming to ultimately integrate the two datasets, we assessed on WH datasets the impact of not using FA as an additional intensity feature for BIANCA. We compared the FA inclusion/exclusion cases in terms of BIANCA performance, relationship between WMHs and age, and the weight of the scanner variable in the multivariate regression model (see *Evaluation metrics* for details). For testing this option, we only used bias field corrected images and fixed BIANCA training set to be mixed (i.e. including examples from WH1 and WH2).

#### Retrospective harmonisation of Whitehall and UK Biobank datasets

We then extended the investigation to include data from the UKB cohort. In this case, no prospective harmonisation had been performed for imaging or non-imaging variables. The cohorts, despite being aging populations, differ in many aspects (see Table 1 for details). Hence, both non-imaging and imaging data required harmonisation.

- **Non-imaging harmonisation**: non-imaging data available for both WH and UKB were converted to a common format. The conversion was conducted using the FMRIB UKBiobank Normalisation, Parsing And Cleaning Kit (FUNPACK https://git.fmrib.ox.ac.uk/fsl/funpack/), a Python library for pre-processing of UKB data containing a large number of procedures allowing us to perform various data sanitisation and processing steps. We defined a configuration file for FUNPACK, currently available online on GitLab (https://issues.dpuk.org/eugeneduff/wmh_harmonisation). It includes both built-in rules and new conversion functions that allowed us to obtain non-imaging variables expressed in the same units of measurements.
- **Imaging data harmonisation – effect of training set composition for BIANCA:**for WH-UKB integration, the manual WMH masks were generated by different raters, bias field correction was already performed as part of the automated pre-processing pipeline (Alfaro-Almagro et al., 2018) and FA was not used as additional intensity feature. We therefore tested whether the use of a specific training set for BIANCA could improve harmonisation between UKB and WH, despite different raters providing WMH examples and the use of only T1 and FLAIR as intensity features. As in the previous scenario, we compared the impact of site-specific and mixed training sets (this time combining examples from WH1, WH2 and UBK). Also in this case, the evaluation included comparing BIANCA performance, the relationship between WMHs and age, and the weight of the scanner variable in the multivariate regression model (see *Evaluation metrics* for details).

#### Evaluation metrics

We evaluated the success of harmonisation in several ways.

First, the harmonised WMH segmentation pipeline should have the same (or as close as possible) WMH segmentation performance across datasets. To assess this, we calculated a series of overlap measures: Dice Similarity Index (DI), voxel-level False Positive Ratio (FPR), voxel-level False Negative Ratio (FNR), cluster-level FPR, cluster-level FNR (see (Griffanti et al., 2016) for details) between manual WMH masks and automatically segmented WMH masks (obtained using leave-one-out cross-validation whenever appropriate). We matched the number and the approximate lesion load of the manually annotated scans used to evaluate the automatic segmentation performance for all datasets (12 subjects for each dataset, WH1, WH2, UKB). We then looked at how different these metrics were between datasets for each option tested (*across-scanner evaluation* within option). In the scanner upgrade scenario we compared metrics between SC1 and SC2 for each of the following options: (A) without BC, single-site training, FA included; (B) with BC, single-site training, FA included; (C) with BC, site-specific training, FA included; (D) with BC, mixed training, FA included; (E) with BC, mixed training, FA excluded. For the WH-UKB harmonisation we compared SC1 vs SC2 vs UKB for the (A) site-specific training and (B) mixed training options (both with BC and no FA).

Alongside the harmonisation aim, we also took into account the accuracy of the WMH segmentation (since consistent BIANCA performance across datasets does not necessarily correspond to accurate performance). Therefore – for each dataset – we compared BIANCA performance across different options ((A) vs (B) for bias field, (B) vs (C) vs (D) for training set, (D) vs (E) for effect of FA – for the scanner upgrade scenario; (A) vs (B) for training set – for the WH-UKB scenario) to investigate whether the adoption of one of them could lead to substantial improvements in terms of either segmentation accuracy, sensitivity or specificity (*within-scanner evaluation* across options).

When the number of available options for both the across- and within-subject factors (being dataset and analysis option, respectively) was equal to two (as for the rater, bias field, and FA assessment) we used two-sample independent t-tests and paired t-tests for statistical assessment. When the number of available options was higher than two (as for the training set assessment) we first performed a two-way mixed ANOVA test, to test for potential interaction between factors and then, if results were significant, we investigated the main effect of each factor through separate one-way ANOVA tests.

We then extended the evaluation to the full sample by considering the output of the automatic WMH segmentation for all the available subjects (WH1=528, WH2=211, UKB=2285), instead of just for those with manual WMH mask. We calculated WMH volumes (expressed as % of total brain volume) and compared them across datasets for each option of the two scenarios. In doing this we wanted to take sources of biological variability into account. Given that age is known to be among the strongest risk factors for WMHs, we started by looking at the correlation between WMH volumes and age in our datasets. We implemented a one-way ANCOVA test, using WMH volumes as the dependent variable, age as the main covariate and scanner/site as the categorical factor. Age was demeaned to avoid multicollinearity and make results more interpretable. With this test we assessed differences in terms of slope (interaction between age and scanner) and intercept at mean age (main effect of scanner) for each option. Similar regression slopes (no significant interaction) and reduced or no volume bias (no significant main effect of scanner) would indicate successful harmonisation.

Finally, harmonisation was evaluated by the extent to which it reduced the variation in WMH volumes that could be explained by scanner and dataset. We assessed this by examining the fit of a linear multivariate model, estimated using Elastic Net (Pedregosa et al., 2011), that predicted WMH volumes from non-imaging variables (see Table 1 for details) (including a variable associated with scanner/dataset). Well harmonised datasets will have minimal variance attributed to the scanner/dataset variables.

## RESULTS

### Scanner upgrade (Whitehall)

#### Effect of rater

Overall, BIANCA produced more consistent WMH masks than manual operators (Fig. 1). Comparing manual and automatic segmentation procedures in terms of between-rater variability (R1 vs R2), we obtained opposite results when considering either the first (R2a) or second rating (R2b) from the second rater. The comparison between R1 and R2a highlighted a higher agreement (higher DI values) between manual masks (M1 vs M2a) than between the corresponding BIANCA output (B1 vs B2a) (Fig. 1.A, p<0.001 paired t-test). On the other hand, the comparison between R1 and R2b showed better consistency for BIANCA results (B1 vs B2b) than manual annotations (M1 vs M2b) (Fig. 1.B, p<0.001 paired t-test). It is worth noting that the worst agreements (both between manual masks and BIANCA outputs) were observed for subjects characterised by very low WMH loads (dotted lines). For within-rater (R2) variability, we observed that BIANCA outputs (B2a vs B2b) had higher consistency than manual masks annotated twice by the same operator (M2a vs M2b) (Fig. 1.C, p<0.001 paired t-test). For details refer to Supplementary Table S1.

**Figure 1.**
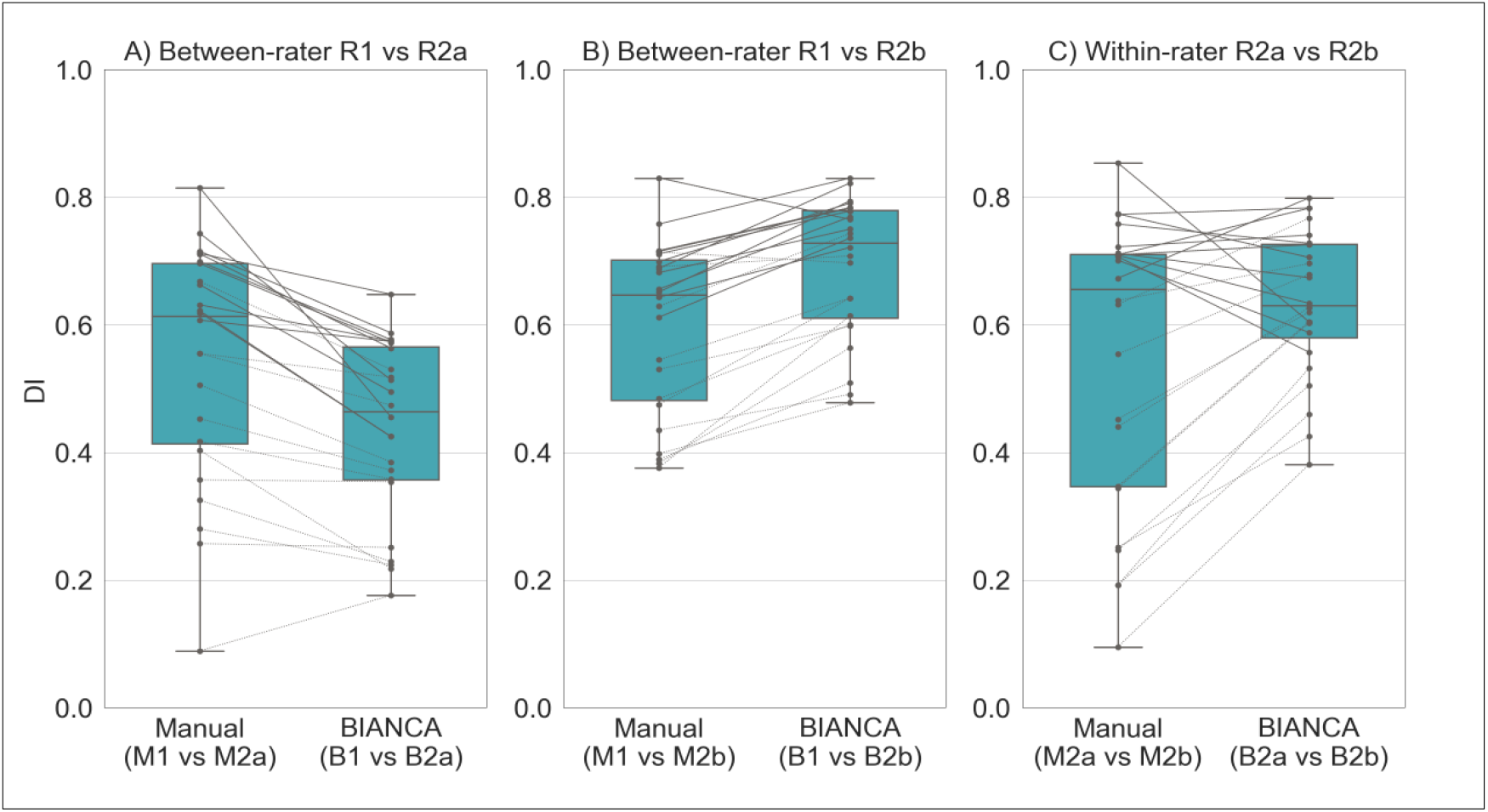
Effect of rater, assessed both in terms of between-(A and B) and within-rater variability (C). Each panel displays a comparison of the agreement (measured with Dice Similarity Index) between manual masks annotated by the raters (left boxplots) and BIANCA outputs generated with masks from those raters (right boxplot). Solid and dotted lines refer to results obtained on subjects characterised, respectively, by high and low WMH load. Legend: R1 = rater 1, R2a = Rater 2, first rating, R2b = rater 2, second rating (1 year apart from the first rating, blind to first rating), M = manual, B = BIANCA.

#### Effect of bias field correction

Bias field correction (BC) led to increased image similarity, when comparing ‘traveling heads’ data from the two WH scanners, as clearly visible from the example shown in Fig. 2.A. This was confirmed by a significant decrease in the cost function (correlation-ratio) after BC (p<0.001 paired t-test; Fig. 2.B).

**Figure 2.**
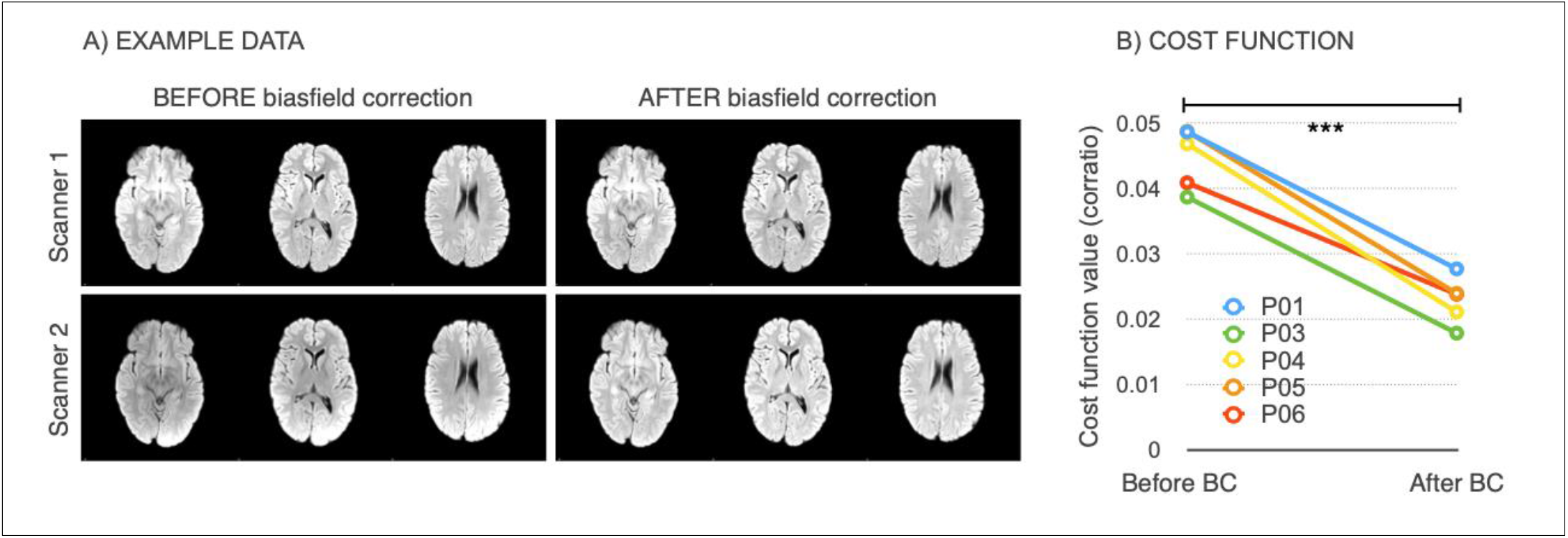
Effect of bias field correction (BC) on ‘travelling heads’ data from the WH dataset. (A) example data from 1 subject acquired on both scanners, before and after BC showing improvement in image similarity after BC (B) Cost function (correlation ratio) between Scanner1/Scanner2 images of the 5 traveling head participants, calculated before and after BC (*** −p < 0.001).

The effects of bias field correction on BIANCA performance are shown in Table 3 and Fig. 3 (where Fig. 3 displays DI values, while the equivalent plots for the other metrics are reported in the Supplementary material). Comparing segmentation performance within scanner, we observed a significant increase in the overall segmentation accuracy after BC, with higher DI values for both datasets (Fig. 3). Moreover, BC led to a greater level of specificity for the WH2 dataset, demonstrated by a significant decrease in FPR and cluster-level FPR. For the WH1 dataset, the DI improvement was accompanied by a decrease of FNR and cluster-level FNR values. This was at the expense of an increase in the WH1 FPR and cluster-level FPR. There was a significant difference in DI values between WH1 and WH2 after but not before BC suggesting worse comparability after BC. However, BC also had a positive impact on FPR which were no longer significantly different across-scanners.

**Table 3.**
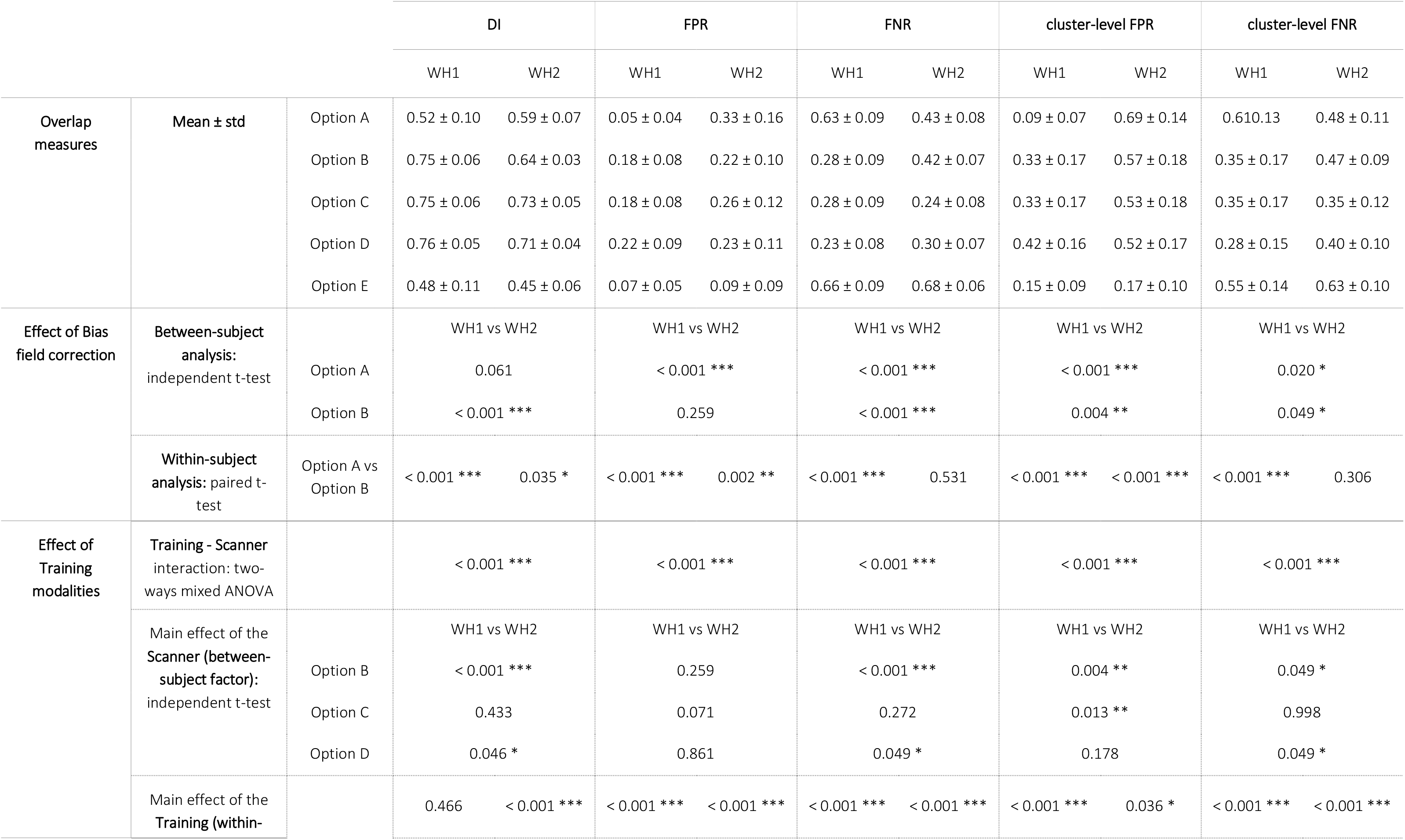
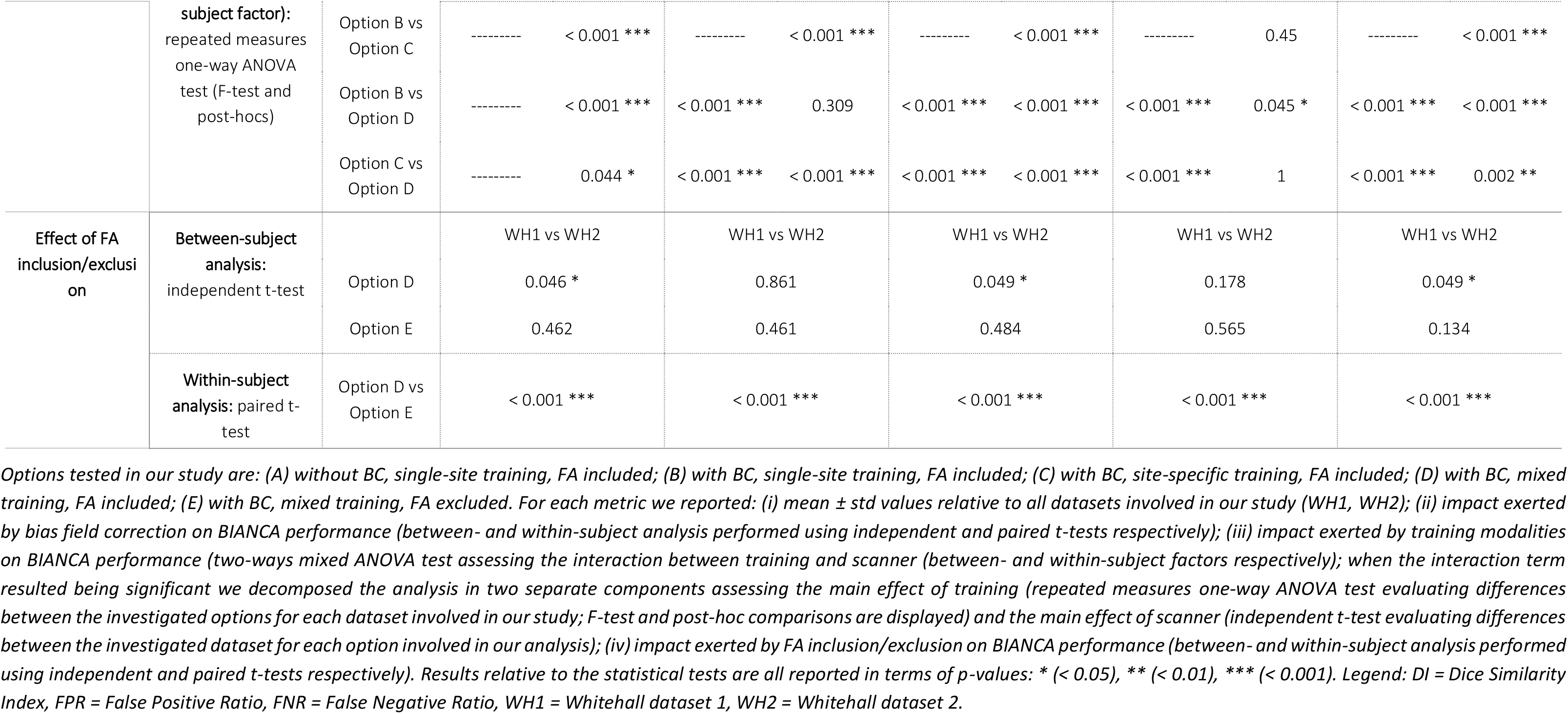
BIANCA performance – scanner upgrade scenario – Summary of all the overlap measures between BIANCA output and the corresponding manual mask, calculated for the different analysis options tested in our study (using leave-one-out cross-validation whenever appropriate). Statistical tests performed on data to assess the impact of bias field correction, training modalities and FA inclusion/exclusion on the segmentation performance.

**Figure 3.**
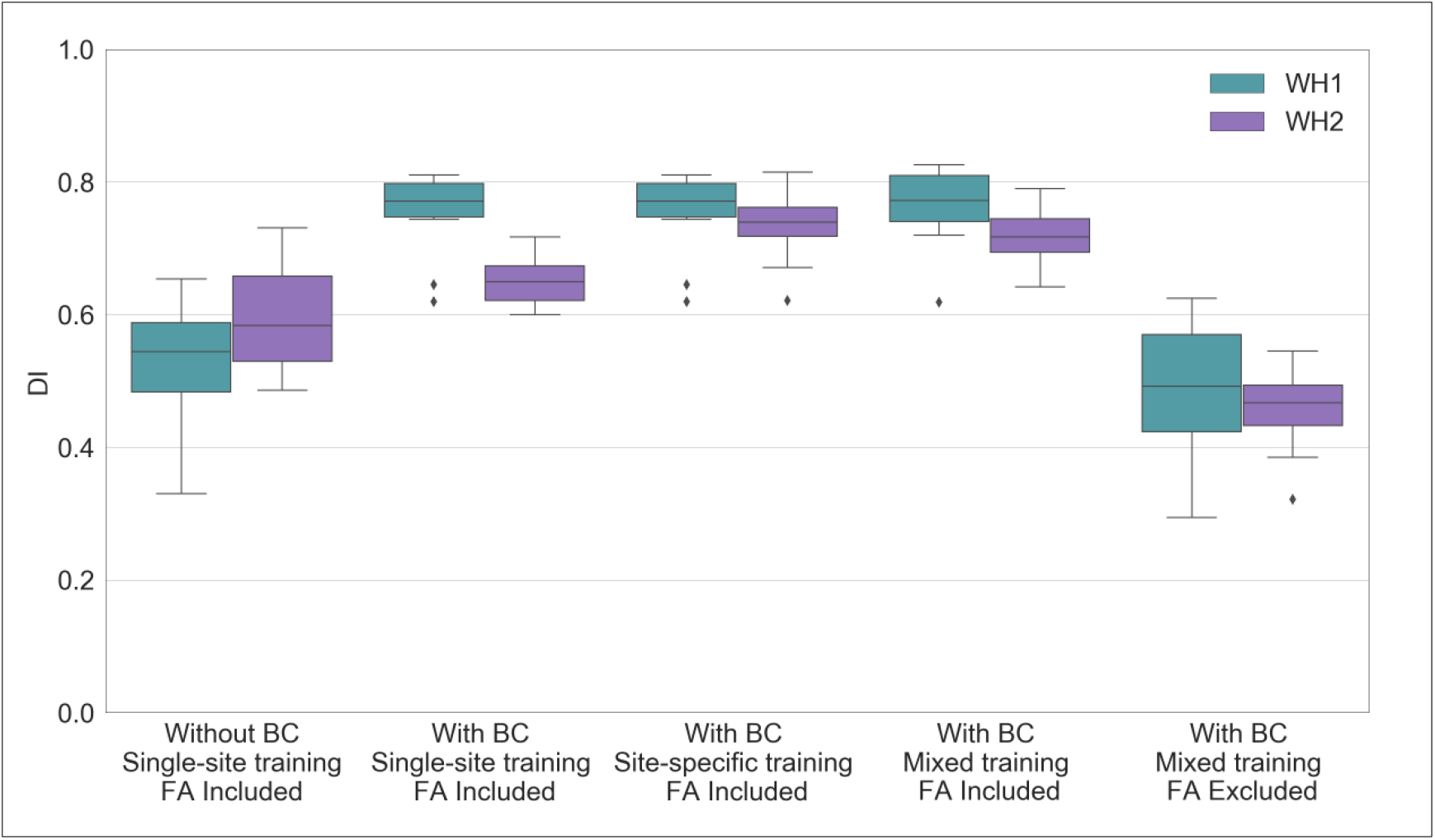
BIANCA performance – scanner upgrade scenario. Box-plot of the Dice Similarity Index (DI) between BIANCA output and the corresponding manual masks for the different analysis options tested during our study (specified on the x axis). All the displayed results were evaluated on a sub-sample of manually segmented subjects (12 for WH1 and 12 for WH2) balanced in terms of WMH load and using leave-one-out cross-validation whenever appropriate (details in the main text).

We then analysed the correlation between WMH volumes and age to determine the extent to which this relationship was affected by the scanner for the two BC options (Fig. 4.A and 4.B). Results of the one-way ANCOVA tests reported in Table 5 show no significant difference when comparing regressions slopes between scanners for both options (p-value=0.782 before BC; p-value=0.789 after BC). A significant across-scanner difference was instead found in the intercepts – in correspondence of the mean age – both before and after BC. However, the difference was reduced after BC (p-value<0.001 before BC; p-value=0.023 after BC).

**Figure 4.**
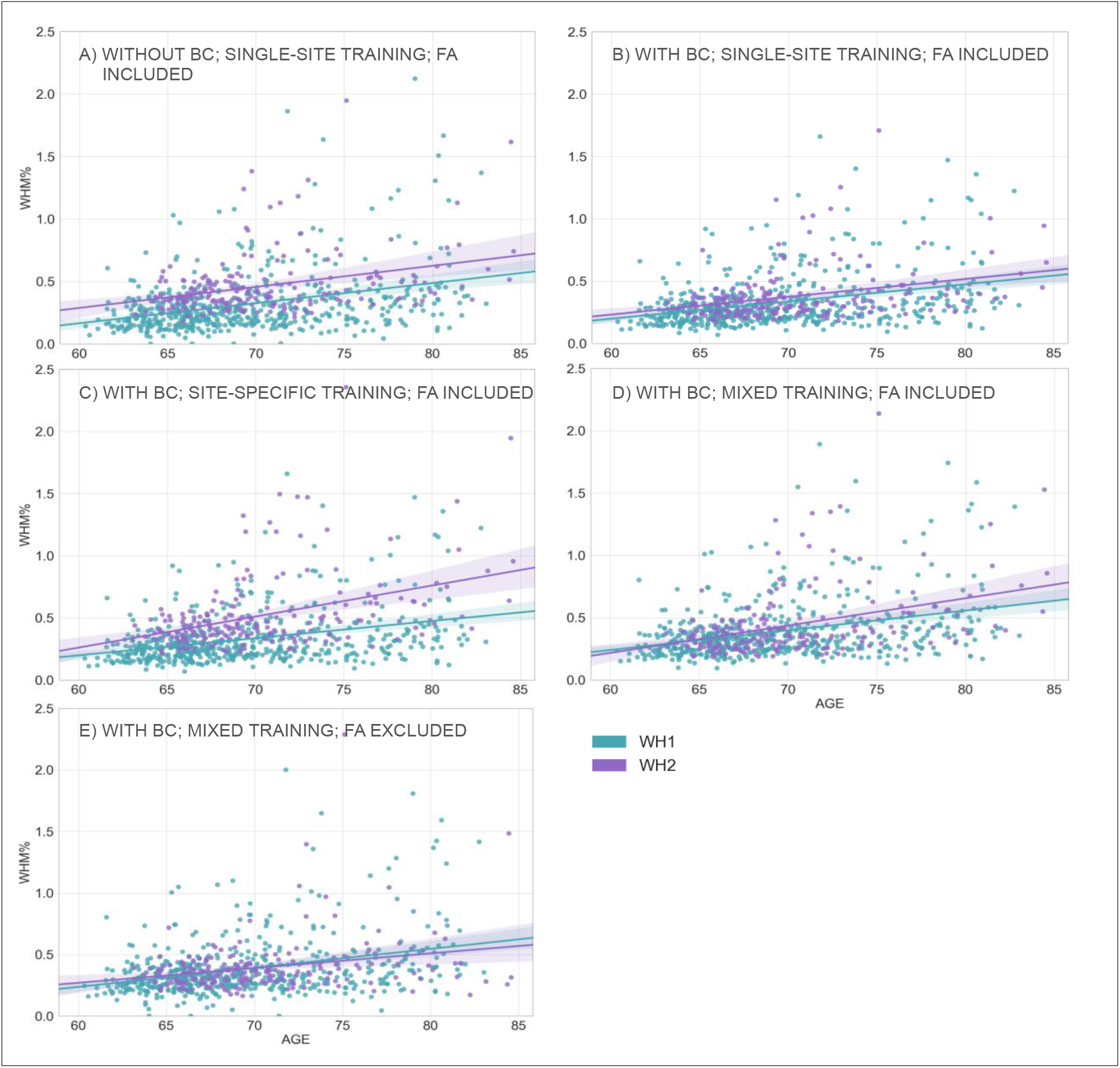
Association between WMHs and age – scanner upgrade scenario. Scatter plot of the relationship between WMH volumes (expressed as % of total brain volume, y axis) and age (x axis), for WH1 (cyan) and WH2 (purple) data. Regression lines with 95% confidence interval are also displayed. Each plot refers to one of the investigated analysis options: (A) without BC, single-site training, FA included; (B) with BC, single-site training, FA included; (C) with BC, site-specific training, FA included; (D) with BC, mixed training, FA included; (E) with BC, mixed training, FA excluded. Evaluation was conducted on the full sample of data for both datasets (WH1 = 528, WH2 = 211).

**Table 4.**
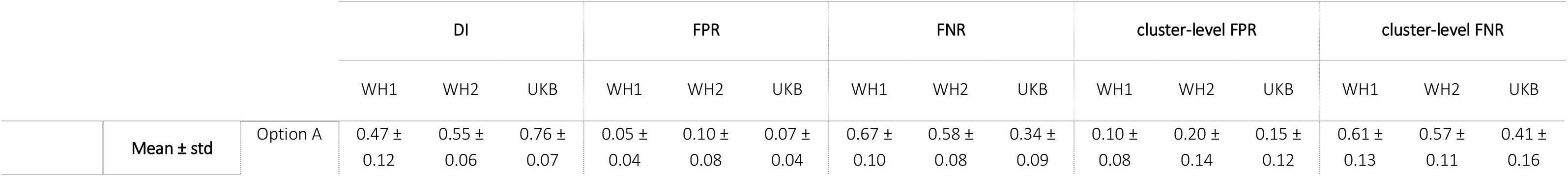
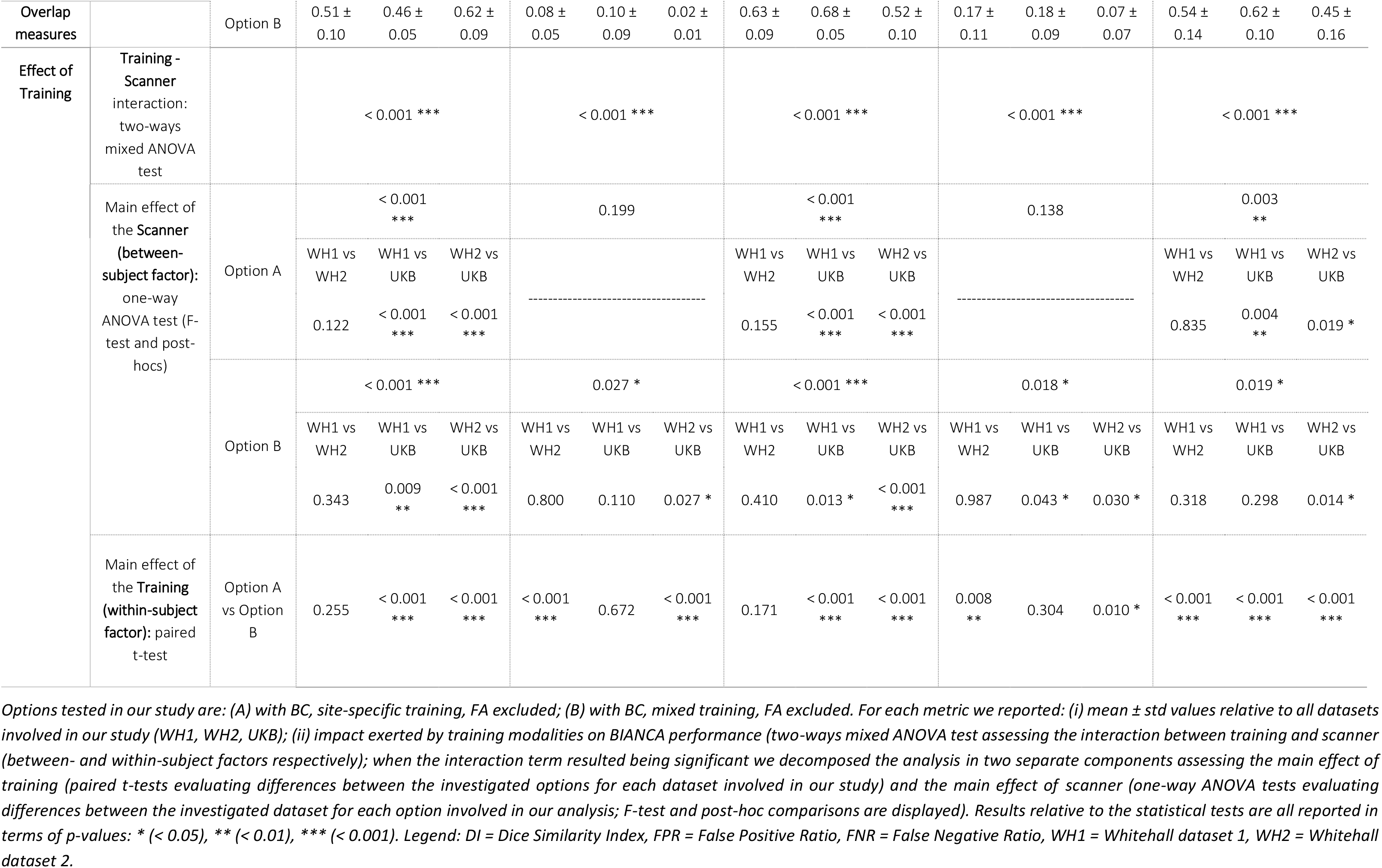
BIANCA performance – retrospective scenario – Summary of all the overlap measures between BIANCA output and the corresponding manual mask, calculated for the different analysis options tested in our study (using leave-one-out cross-validation). Statistical tests performed on data to assess the impact of training modalities on the segmentation performance.

**Table 5.**
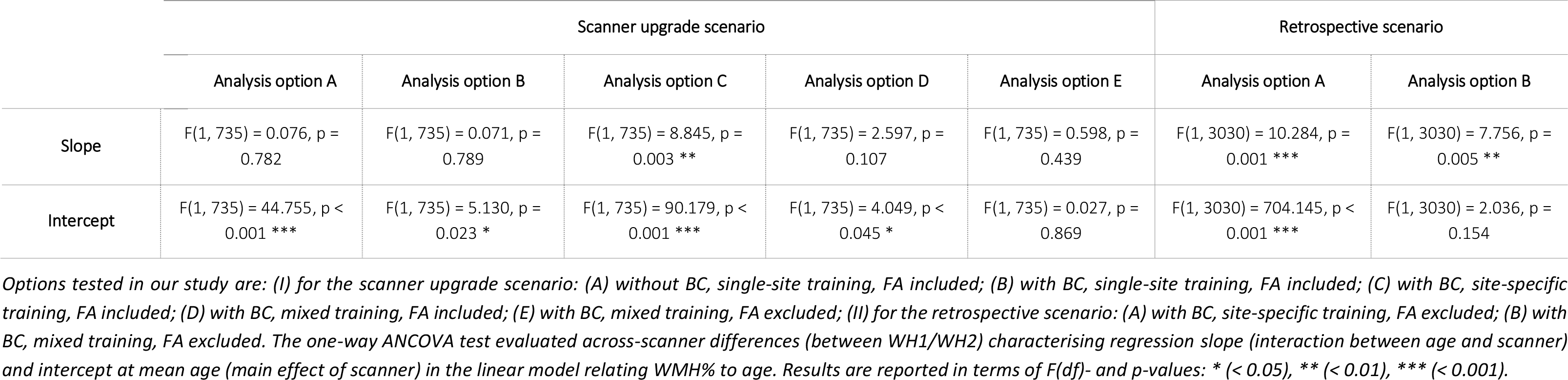
Analysis of the relationship between WMH volumes and age – scanner upgrade and retrospective scenario – Summary of the one-way ANCOVA test.

Finally, the implemented Elastic Net model showed that, after BC, the amount of variance in WMH volume attributed to the scanner/site of acquisition was lower, passing from second to sixth position (Fig. 5.A and 5.B, and Table 6 for specific values).

**Figure 5.**
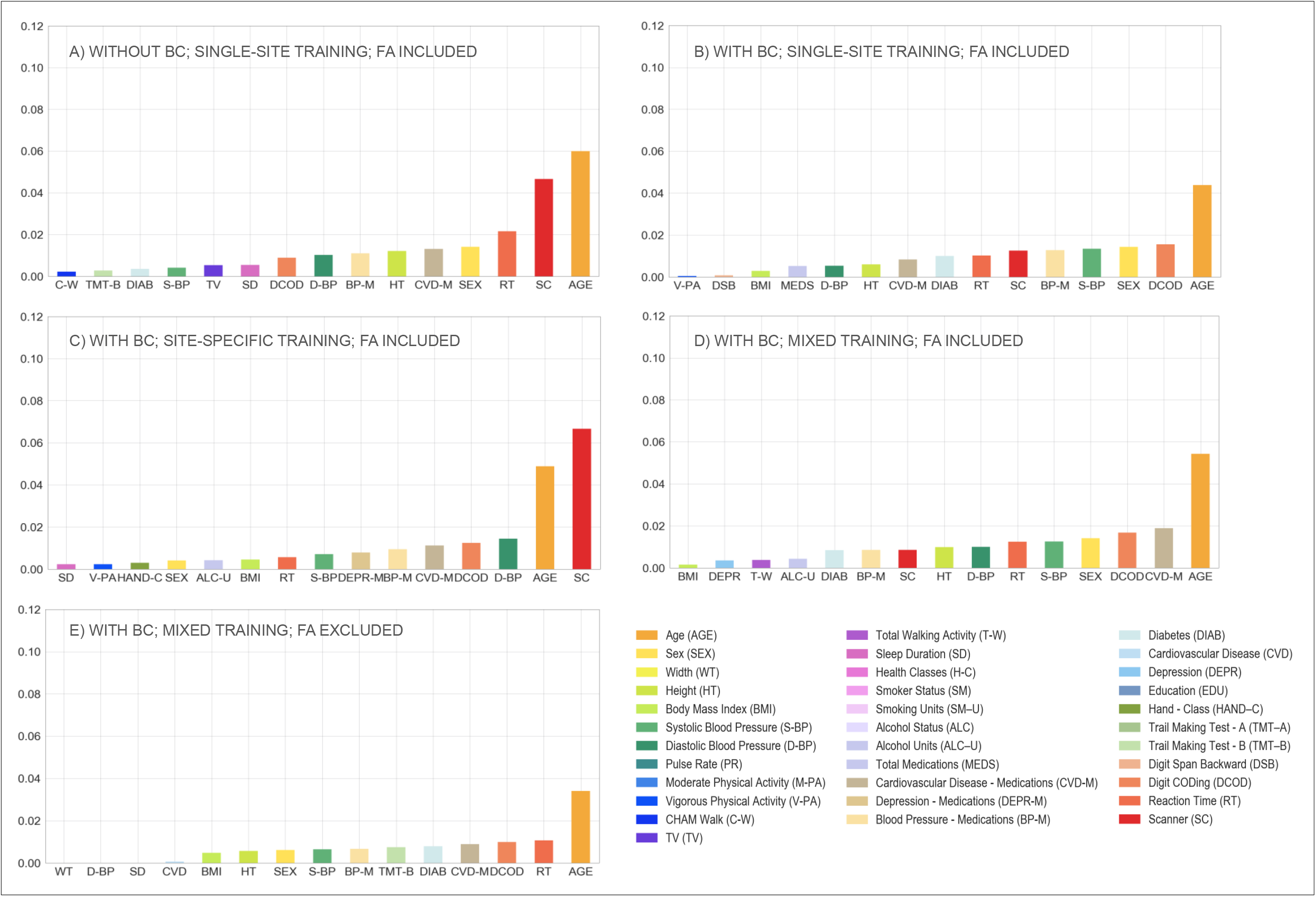
Multivariate model – scanner upgrade scenario. Percentage of variance (y axis) explained by non-imaging variables (reported on the x axis) in the linear multivariate model that was implemented (Elastic Net). Evaluation was conducted on the full sample of data (WH1 = 528, WH2 = 211). Each plot refers to one of the investigated analysis options: (A) without BC, single-site training, FA included; (B) with BC, single-site training, FA included; (C) with BC, site-specific training, FA included; (D) with BC, mixed training, FA included; (E) with BC, mixed training, FA excluded. Variable scanner/site (SC) highlighted in red. Values are reported in Table 6 and supplementary table S2.

**Table 6.**
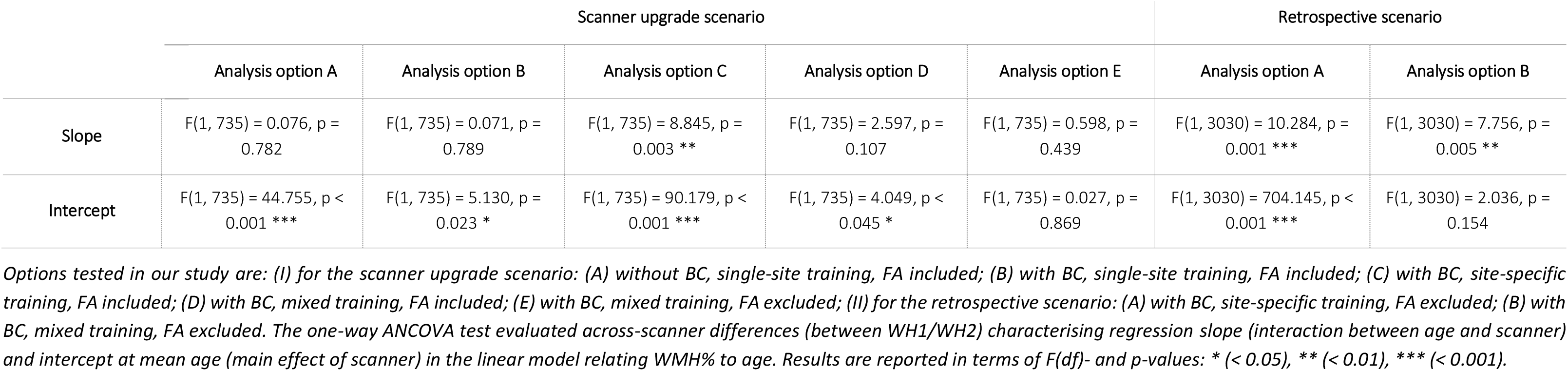
Elastic Net Regression performance – scanner upgrade and retrospective scenario – Summary of the results in terms of variance explained by the model and by a subset of the features which constituted it. Variables scanner and age are here reported for the different analysis options tested in our study. Full details of all the other features are provided in supplementary table S2.

#### Effect of training set composition for BIANCA

Overall, our results suggest that the mixed-training option offers the best trade-off among the explored evaluation metrics, providing good and consistent BIANCA performance and consistent WMH volumes.

When investigating the presence of a significant interaction between scanners (WH1/WH2) and training options (single-site/site-specific/mixed), a two-way mixed ANOVA test gave significant results for all the assessed overlap measures (Table 3). Therefore, we investigated the effect of each factor separately, evaluating firstly across-scanner and then within-scanner performances. Site-specific training produced the most consistent segmentations with respect to across-scanner performance. Between the remaining two options, the mixed training showed better consistency with respect to single-site training, with no significant WH1-WH2 difference in the cluster-level FPR values. When comparing segmentation performance within-scanner, we observed an overall improvement of results from single-site to site-specific training for WH2 (for WH1 is the same option as both the single-site and site-specific training represent 24 subjects from WH1 in this case). The significant improvements in DI, FNR and cluster-level FNR were at the only expense of increased FP values. The comparison between site-specific and mixed training led to different results for the two scanners, with significantly worse FPR and cluster-level FPR values for mixed training in WH1 and worse DI, FNR and cluster-level FNR values in WH2. The remaining indicators showed improved or unaltered performance with mixed training. When comparing single-site and mixed training the results showed a favourable pattern towards the latter. For WH1 we observed a significant improvement for both FNR and cluster-level FNR when using mixed training, no significant difference in DI, and worse FPR and cluster-level FPR. For WH2, better performances were observed using a mixed training for all the indicators except FPR, which was not significantly different from the single-site training case.

The results obtained from the one-way ANCOVA tests (Table 5) showed that site-specific training led to a significant difference between the age regression slopes for the two scanners (p-value=0.003). Using this option also led to the highest volume bias (calculated at the mean age) between scanners (Fig. 4.B, 4.C and 4.D). The adoption of a mixed training had a positive impact on regression slopes, such that they were no longer significantly different (p-value=0.107) and also reduced the volume bias (at the mean age) – although it was still significant (p-value=0.045).

When site-specific training was used, the weight of the scanner/site variable was greatly increased in the multivariate regression model, compared to the single-site option, with scanner/site being the variable that explained the greatest amount of variance (Fig. 5.B and 5.C, and Table 6 for specific values). The adoption of a mixed training instead, reduced the amount of variance explained by the scanner/site variable, with the variable moving to the ninth position (Fig. 5.D, and Table 6 for specific values).

#### Effect of FA information

The removal of FA as an additional intensity feature for WMH segmentation led to higher consistency between sites, but lower segmentation accuracy.

Without FA there were no significant differences between the WH1 and WH2 datasets in all performance metrics. There was a significant decrease in the overall segmentation accuracy when excluding FA from the intensity features used by BIANCA, with lower DI performances (Fig. 3), and a negative impact on both FNR and cluster-level FNR (worse sensitivity). Removing FA also lowered FPR and cluster-level FPR, leading to a greater level of specificity.

For the correlation between WMH volumes and age, results of the one-way ANCOVA tests (Table 5) showed that, excluding FA, the difference in slopes remained not significant (p-value=0.439). The already small volume bias (at mean age) was further decreased (Fig. 4.D and 4.E) and, indeed, the difference between intercepts was no longer significant (p-value=0.869).

Extracting WMHs using FLAIR and T1-weighted images only led to a decrease in the variance explained by the scanner/site variable, which was no longer present amongst the most predictive features (Fig. 5.D and 5.E, and Table 6 for specific values).

### Retrospective harmonisation of Whitehall and UK Biobank datasets

#### Non-imaging harmonisation

By applying our configuration file for FUNPACK, we brought all the variables into the same units for both datasets. Table 1 shows the format/units that each of the selected non-imaging variables were originally acquired with in WH and UKB, as well as the harmonised units chosen and the resulting harmonised mean and standard deviation values.

#### Imaging data harmonisation – effect of training set composition for BIANCA

We next assessed the impact of different training sets (site-specific and mixed training) on the level of harmonisation between the WH and UKB WMH datasets (the single-site training was not tested, as it gave the worst results in the scanner upgrade scenario).

Results, relative to BIANCA performance, in terms of Dice Similarity Index (DI) are shown in Fig. 6. The equivalent plots for the other metrics are reported in the Supplementary material. The two-way mixed ANOVA test highlighted the presence of a significant interaction between the scanners (WH1/WH2/UKB) and the training options (site-specific/mixed) for all the overlap measures (Table 4). For this reason, we further evaluated the main effect of each factor, investigating across- and within-scanner performance separately. Results of the one-way ANOVA test revealed significant differences for all metrics across scanners when using a mixed training. The site-specific training gave more homogeneous results, (non-significant FPR and cluster-level FPR). Post-hoc pairwise comparisons revealed no significant difference in any overlap metrics between WH1 and WH2 for either of the training options. On the other hand, UKB showed a different performance with respect to the other datasets (WH1, WH2), using either site-specific or mixed training. Significant differences between WH1 and UKB were observed in DI, FNR and cluster-level FNR in the site-specific training case. DI, FNR and cluster-level FPR were significantly different in the mixed training case. All overlap metrics were significantly different between WH2 and UKB, except FPR and cluster-level FPR using site-specific training. Within-scanner comparisons highlighted a more favourable pattern towards the site-specific training. In fact, the use of a mixed training dataset led to improved segmentation sensitivity only for WH1, with a significant decrease of cluster-level FNR, and improved specificity for UKB with lower FPR and cluster-level FPR.

**Figure 6.**
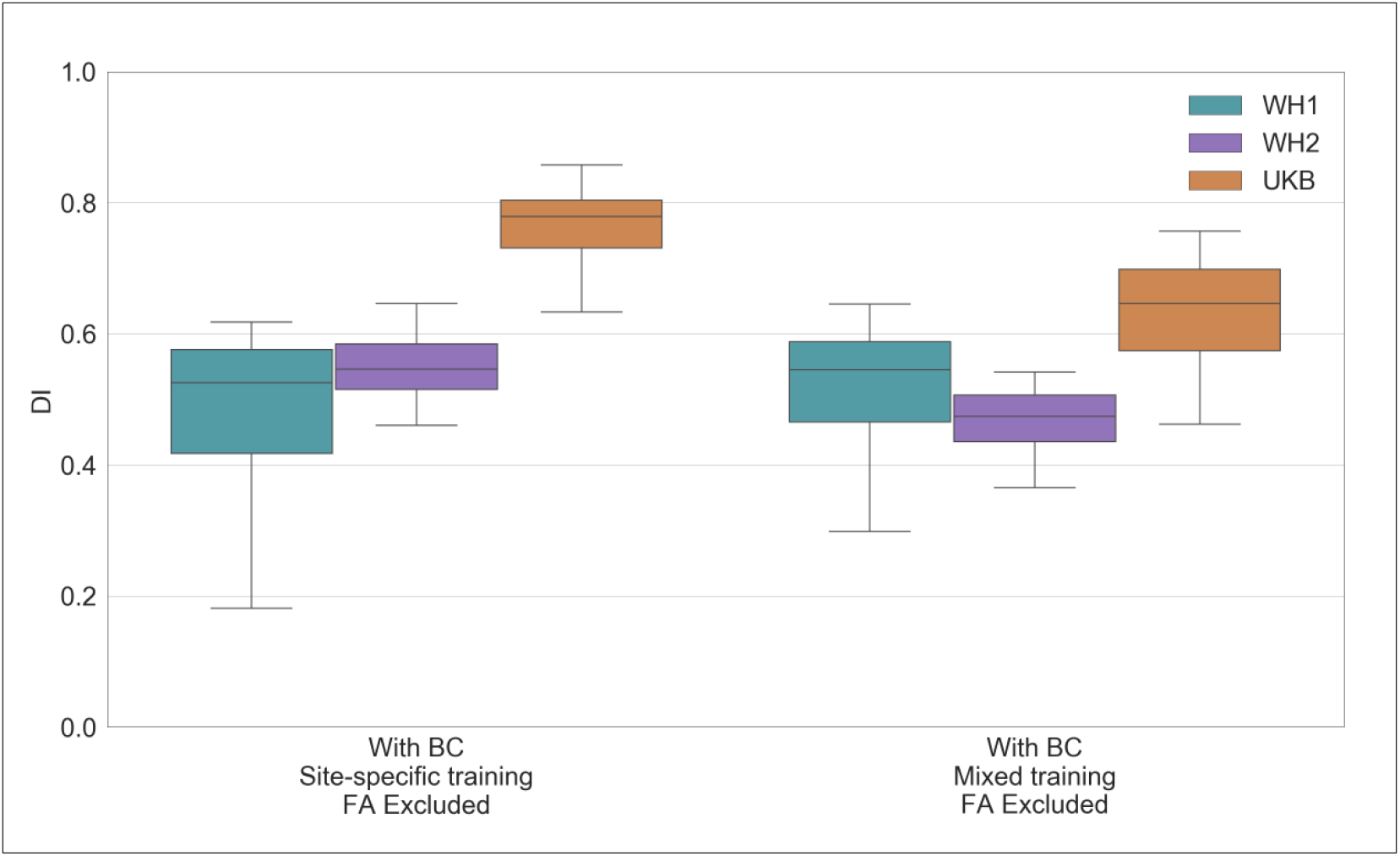
BIANCA performance – retrospective data merging scenario. Box-plot of the Dice Similarity Index (DI) between BIANCA output and the corresponding manual mask for the different analysis options tested during our study (specified on the x axis) All the displayed results were evaluated on a sub-sample of manually segmented subjects (12 for WH1, 12 for WH2 and 12 for UKB) balanced in terms of WMH load and using leave-one-out cross-validation.

In terms of correlation between WMH volumes and age, we compared results for WH1, WH2 and UKB (Table 5). With respect to the site-specific case, the adoption of a mixed training led to a decrease in the across-scanner difference in regression slopes, even if it still remained significant (p-value=0.005 mixed training; p-value=0.001 site-specific training, one-way ANCOVA test). The significant bias (evaluated at the mean age) characterising the site-specific case was substantially decreased using a mixed training set (Fig. 7.A and 7.B). Indeed, the difference in regression intercepts at mean age was no longer significant (p-value=0.157, one-way ANCOVA test).

**Figure 7.**
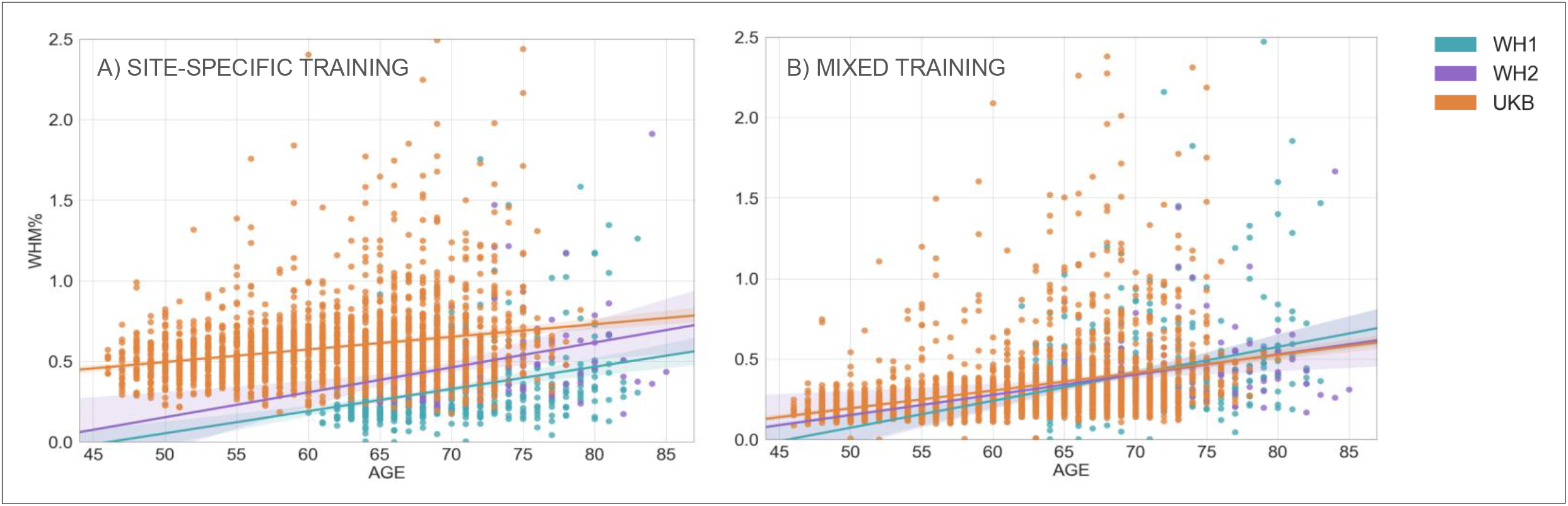
Association between WMHs and age – retrospective data merging scenario. Scatter plot of the relationship between WMH volumes (expressed as % of total brain volume, y axis) and age (x axis), for WH1 (cyan), WH2 (purple) and UKB (orange) data. Regression lines with 95% confidence interval are also displayed. Each plot refers to one of the investigated analysis options: (A) with BC, site-specific training, FA excluded; (B) with BC, mixed training, FA excluded. Evaluation was conducted on the full sample of data for all datasets (WH1 = 528, WH2 = 211, UKB = 2295).

The Elastic Net model showed that the scanner/site was no longer present amongst the most predictive features when using mixed training, compared to site-specific training where it explained the highest amount of variance in WMH volumes (Fig. 8.A and 8.B, and Table 6 for specific values).

**Figure 8.**
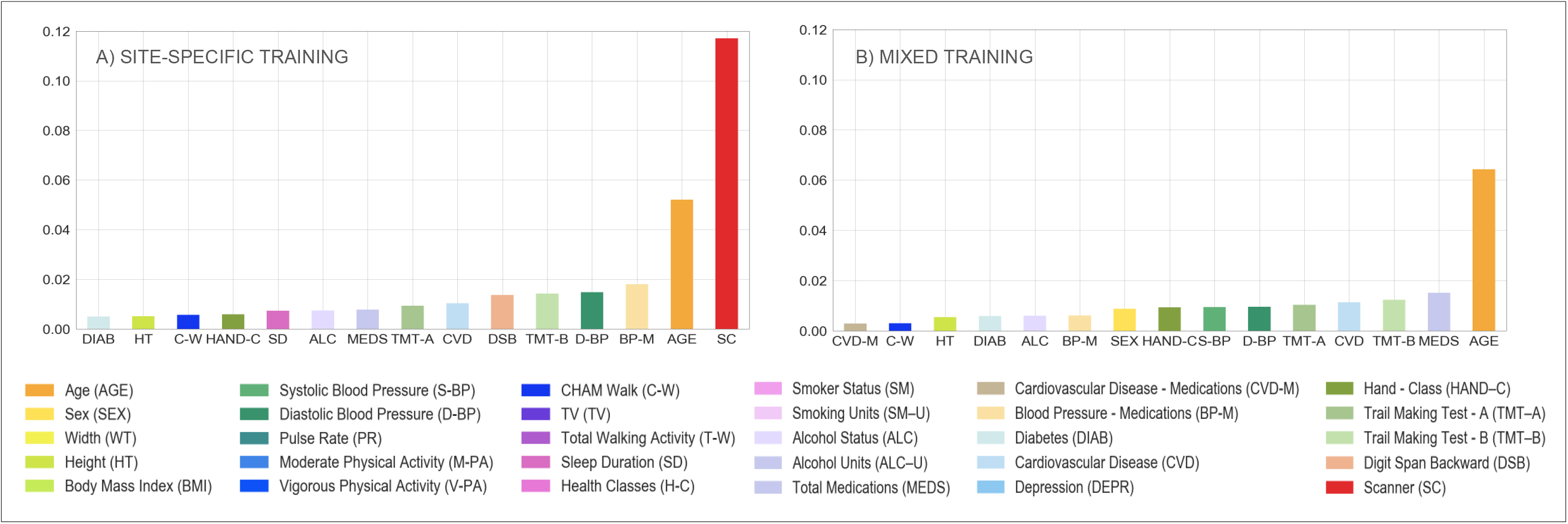
Multivariate model – retrospective data merging scenario. Percentage of variance (reported on the y axis) explained by non-imaging variables (reported on the x axis) in the linear multivariate model that was implemented (Elastic Net). Evaluation was conducted on the full sample of data for all the involved populations (WH1 = 528, WH2 = 211, UKB = 2295). Each plot refers to one of the investigated analysis options: (A) with BC, site-specific training, FA excluded; (B) with BC, mixed training, FA excluded. Variable scanner/site (SC) highlighted in red. Values are reported in Table 6 and supplementary table S2.

## DISCUSSION

In this work we present an analysis of the harmonisation of measures of white matter hyperintensities (WMHs) of presumed vascular origin across different large-scale datasets. We dealt with data from three scanners across two studies on healthy ageing. The study design allowed us to assess two different scenarios: a scanner upgrade (analogous scenario to a multi-centre study, involving a single population acquired with the same acquisition protocol on two MRI scanners) and a retrospective data merging (two distinct large populations acquired with different acquisition protocols on different MRI scanners). Each dataset included both imaging and non-imaging data that were exploited to develop harmonisation strategies and evaluate the results. We used an automated segmentation tool, BIANCA, to extract WMH measures from each imaging dataset and investigated the impact of different factors on the comparability of WMH measures: the rater performing manual segmentation of the examples used to train BIANCA, the process of bias field correction of the FLAIR images, the composition of the dataset used to train BIANCA (training set) and the inclusion/exclusion of FA as one of the MRI modalities. We investigated different processing strategies aiming to find the combination that led to the most consistent results across scanners or studies. We evaluated the success of each strategy looking for the best trade-off between consistency and accuracy of segmentation performance, and consistency of WMH volumes, after modelling the biological variability in the datasets (age and other non-imaging variables related to WMHs).

BIANCA needs to be trained by providing manual WMH segmentations, which are known to be affected by inter- and intra-rater variability (Guo et al., 2019). We wanted to assess how BIANCA would cope with this source of variability. To this aim we tested if BIANCA trained with different manual masks (either multiple annotations by different raters or repeated annotations by the same rater) would produce WMH masks that are more or less consistent than manual masks themselves. On data from a single scanner (WH1) we observed that the consistency of the manual segmentation of the data has a major impact on the final BIANCA outputs. If the manual segmentations provided to BIANCA are sufficiently similar between raters/ratings, the automated tool improves the consistency of the output, providing better within- and between-rater agreement than the manual raters/ratings themselves. On the other hand, if the agreement between manual masks is low, BIANCA results can be even less consistent than manual masks. This prompts the need to standardise the definition of WMHs, especially in light of the fact that even if an increase in rating consistency is eventually achieved, this does not necessarily mean the obtained results are better in terms of accuracy. While for other segmentation tasks, e.g. hippocampus segmentation, clear protocols exist for manual labelling (Zandifar et al., 2018), there is no such protocol for WMHs. It is also worth noting that the lowest agreements (both between manual and automatic results) were observed for subjects characterised by a very low WMH load. In these images, WMHs are likely to be more difficult to segment because of their less obvious appearance or small size. Specific guidelines should therefore aim to clarify these sources of ambiguity. This analysis was limited by the relatively small number of ratings available and the range of expertise of the raters (R1 neuroimaging researcher, R2 medical student trained and supervised by an experienced neurologist). However, the scope of this evaluation was to explore how differences in manual ratings can impact a supervised segmentation method like BIANCA. To help quantifying the variability caused by manual segmentation we looked at the average agreement (DI) range and found that our between- and within-rater agreement is comparable with the scan-rescan agreement in WMHs assessed in a previous study (inter-scanner range: 0.63–0.65; intra-scanner range: 0.63–0.77) (Guo et al., 2019). This suggests that the impact of the rater on the final segmentation is comparable to the effect of repeating the acquisition using the same settings.

Correcting for bias field had a positive impact on almost all the metrics used for evaluation, indicating that, overall, its adoption is crucial to successful harmonisation. We observed increased image similarity when comparing ‘traveling heads’ data from the WH scanners, showing a clear removal of scanner-related variability in the images. BIANCA performance improved after BC, although in terms of consistency of performance between scanners, an improvement was only observable when BC was combined with a different strategy for the composition of the training dataset, such as BIANCA re-training within each scanner or the merging of multiple examples from different scanners (Fig. 3). The successful removal of non-biological differences with BC was also evident when considering the correlation between WMH volumes and age, which showed that BC preserved the relationship with age (slopes not significantly different) while causing a decrease in the volume bias in correspondence of the mean age. The Elastic Net model confirmed the improved harmonisation with a significant decrease in the importance attributed to the scanner/site of acquisition. Bias field correction of T2-weighted (and FLAIR) images is, however, not always included in pre-processing pipelines. In this work we specifically assessed the impact of BC on WMH segmentation and confirmed that it is beneficial to obtain more consistent image segmentation outputs across datasets.

The information provided by dMRI proved to be useful to obtain accurate WMH segmentation. When using FA maps as one of the intensity features for BIANCA, the performance within-scanner was higher than when using only T1-weighted images and FLAIR. However, when using only two modalities, all the overlap measures were more consistent across scanners, there were no significant differences in the slope of the regression lines of WMH volumes and age, and no significant volume bias (not significant difference in the intercepts at the mean age). Furthermore, the scanner was no longer a significant predictor of WMH volumes in the Elastic Net model. The decision on whether to use FA would therefore depend on the application. While for an accurate segmentation it is useful to include features from diffusion-weighted scans, it also constitutes an additional source of variability across datasets and scanners, leading to less harmonised WMH measures. Harmonisation of diffusion MRI data is currently an active area of research (Fortin et al., 2017; Mirzaalian et al., 2016), and it is likely that this modality will require specific harmonisation strategies that go beyond the scope of this work. Further work in this area will allow integrating DTI-derived measures in multimodal analyses while maintaining good consistency of results. Another aspect to keep in mind is that FA might not always be available (while T2-FLAIR and T1 scans are more commonly acquired), preventing the integration of datasets (or participants within a dataset) that do not have all of them available and usable.

Regarding the choice of the composition of the training dataset for BIANCA we started by exploring three options in the scanner upgrade scenario. We compared the effect of using the same set for all the sites (single site), re-training BIANCA within-scanner (site-specific), or merging examples from different scanners (mixed). The first option led to the biggest difference in BIANCA performance across datasets and a significant bias in the volumes (significantly different intercept at the mean age), although the relationship with age remained consistent (non-significant difference in regression slopes, highest amount of variance explained by age). On the other hand, the second option provided the highest and most consistent BIANCA performance (overlap with manual masks on the subset of subjects with manual labels available) but led to the biggest difference in WMH volumes on the whole sample (significantly different slopes of the regression lines, significantly different intercept at the mean age, highest amount of variance explained by the scanner variable). The results observed for the mixed training set (third option) suggest it represents the best trade-off between good and consistent BIANCA performance, and consistent WMH volumes. Although this could be also due to the fact that more images were used in the mixed training, similar results were observed when using the same number of images (12 from each scanner).

We further compared the best performing options (site-specific vs mixed) when harmonising WMH measures between WH and UKB. Using bias field corrected data and FLAIR and T1 as intensity features, the results were similar to the scanner upgrade scenario. While the segmentation performance was overall higher in the case of site-specific training, the more consistent results were obtained with the mixed training set.

Also in this case, the choice of the most suitable training set should be made depending on the application. When prioritising a more accurate WMH segmentation, a site-specific training is likely to give the best performance. When the aim is to compare or merge multiple datasets, a mixed training set is more appropriate.

It is true that either of the above best options would require the effort of generating, or having access to, some manual masks and having to re-train BIANCA. Even if the numbers required are not high (12 images per dataset proved to be enough), this could still be an unfeasible option for some applications. The use of a single training set for multiple datasets would still be a valid option, but in light of our results, the recommendation would be to carefully check the segmentation accuracy and, when combining the resulting volumes, to consider the use of further strategies in the analyses to address potential biases (e.g. additional covariate in statistical analyses). The fact that including more examples from different datasets improved the results suggests that a promising solution would be to build a larger and more representative/generalisable training set, including examples from more scanners/datasets, that could be widely used. Towards this, we are publicly sharing our mixed training sets. Future work on more datasets should assess if, with a sufficiently large set of examples, a single training set is general enough to be able to be successfully applied to new datasets.

An important part of retrospective data merging was also the harmonisation of non-imaging variables. Modelling the biological variability is crucial to obtain imaging measurements that are well aligned across datasets. The *ad-hoc* configuration file we created for FUNPACK allowed us to obtain matched variables, with the same units across the WH and UKB datasets. The configuration file is openly available at (https://issues.dpuk.org/eugeneduff/wmh_harmonisation). It is fully customizable, so it can be adapted to different datasets and expanded to include more variables and conversion rules.

To conclude, we identified processing strategies to maximise the consistency across two large datasets, Whitehall II and UK Biobank, for the study of WMHs. We harmonised non-imaging variables and proposed a processing pipeline to minimise the effect of non-biological sources of difference in the imaging data. The main recommendations emerging from this work are the following:

- use WMH manual masks generated from the same rater whenever possible and establish guidelines to maximise consistency of the manual masks;
- perform bias field correction;
- use a small set of modalities (T1-weighted and FLAIR), which are more reliably present across studies;
- train BIANCA on data coming from a mix of different scanners/studies when working with more than one dataset.

We showed that with these steps, and appropriate modelling of sample differences, through the alignment of demographic, cognitive and physiological variables, we can provide highly consistent WMH measures. These results open up a wide range of applications for the study of WMHs and potentially other neuroimaging markers across extensive databases of clinical data.

## DATA/CODE AVAILABILITY STATEMENT

The Whitehall study follows MRC data sharing policies (https://www.mrc.ac.uk/research/policies-and-guidance-for-researchers/data-sharing/).

Data will be accessible via the Dementias Platform UK (https://portal.dementiasplatform.uk/) after 2020. This research has been conducted using the UK Biobank Resource (applications number 42596 and 8107).

The FUNPACK configuration file and FSL-BIANCA training sets are openly available (https://issues.dpuk.org/eugeneduff/wmh_harmonisation).

## ETHICS STATEMENT

Whitehall dataset: ethical approval was granted generically for the “Protocol for non-invasive magnetic resonance investigations in healthy volunteers” (MSD/IDREC/2010/P17.2) by the University of Oxford Central University / Medical Science Division Interdisciplinary Research Ethics Committee (CUREC/MSD-IDREC), who also approved the specific protocol: “Predicting MRI abnormalities with longitudinal data of the Whitehall II sub-study” (MSD-IDREC-C1-2011-71).

UK Biobank dataset: informed consent is obtained from all UK Biobank participants; ethical procedures are controlled by a dedicated Ethics and Guidance Council (http://www.ukbiobank.ac.uk/ethics) that has developed with UK Biobank an Ethics and Governance Framework (given in full at http://www.ukbiobank.ac.uk/wp-content/uploads/2011/05/EGF20082.pdf), with IRB approval also obtained from the North West Multi-center Research Ethics Committee.

## ACKNOWLEDGMENTS

We thank all Whitehall II participants for their time, the Whitehall II staff at the University College London, Mandy Pipkin and Barbora Krausova for assisting with recruitment and data collection, the FMRIB Radiographers team for data acquisition, Dr. Christoph Arthofer for the helpful discussions, IT and support teams at the Wellcome Centre for Integrative Neuroimaging for their helpful collaboration.

