## Supplementary Figure 1, 2, 3, 4, 5, 6, 7, 8; Supplementary Table 1, 2. for "Integrating large-scale neuroimaging research datasets: harmonisation of white matter hyperintensity measurements across Whitehall and UK Biobank datasets"

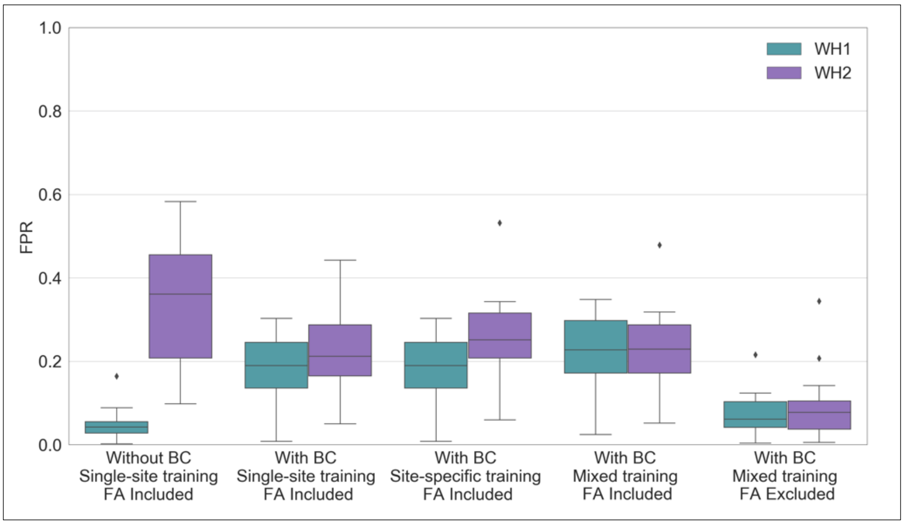


***Figure S1.*** *BIANCA performance – scanner upgrade scenario. Box-plot of the voxel-level False Positive Ratio (FPR) between BIANCA output and the corresponding manual masks for the different analysis options tested during our study (specified on the x axis). All the displayed results were evaluated on a sub-sample of manually segmented subjects (12 for WH1 and 12 for WH2) balanced in terms of WMH load and using leave-one-out cross-validation whenever appropriate.*

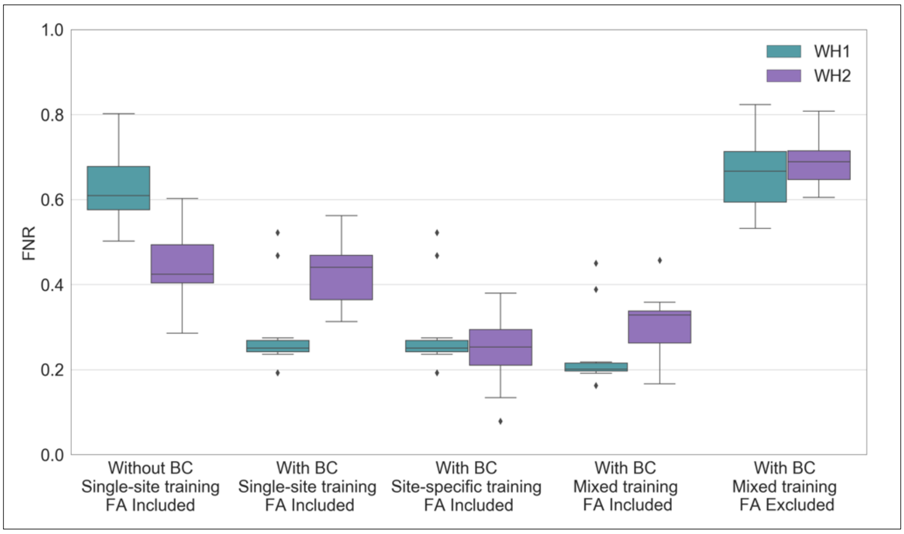


***Figure S2.*** *BIANCA performance – scanner upgrade scenario. Box-plot of the voxel-level False Negative Ratio (FNR) between BIANCA output and the corresponding manual masks for the different analysis options tested during our study (specified on the x axis). All the displayed results were evaluated on a sub-sample of manually segmented subjects (12 for WH1 and 12 for WH2) balanced in terms of WMH load and using leave-one-out cross-validation whenever appropriate.*

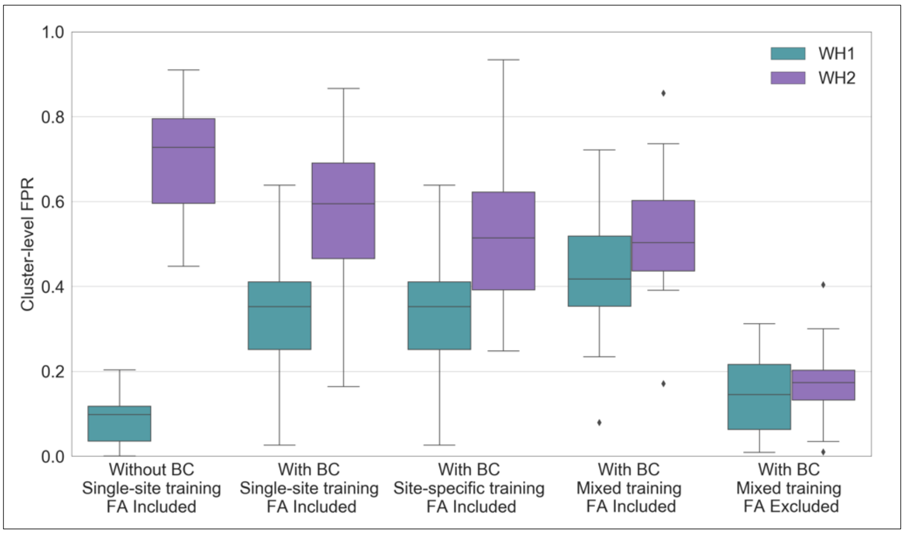

***Figure S3.*** *BIANCA performance – scanner upgrade scenario. Box-plot of the cluster-level FPR between BIANCA output and the corresponding manual masks for the different analysis options tested during our study (specified on the x axis). All the displayed results were evaluated on a sub-sample of manually segmented subjects (12 for WH1 and 12 for WH2) balanced in terms of WMH load and using leave-one-out cross-validation whenever appropriate.*

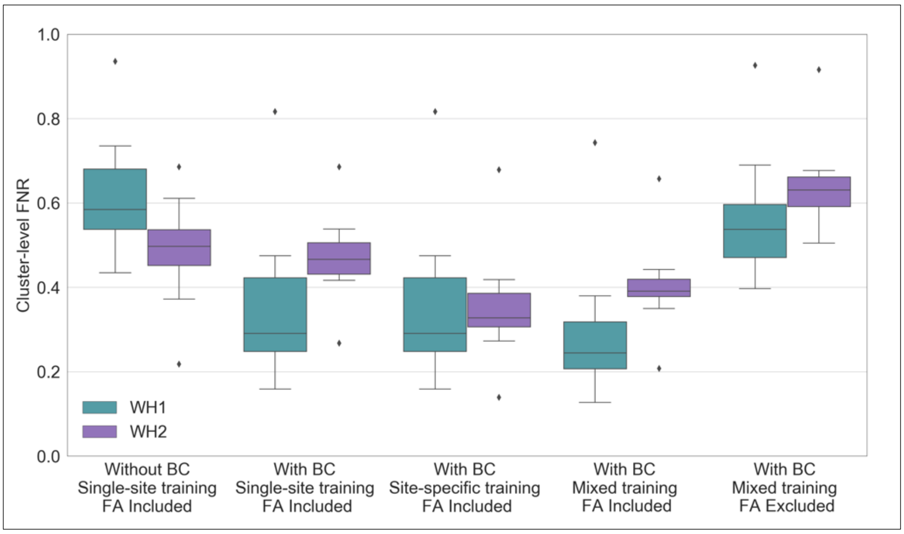


***Figure S4.*** *BIANCA performance – scanner upgrade scenario. Box-plot of the cluster-level FNR between BIANCA output and the corresponding manual masks for the different analysis options tested during our study (specified on the x axis). All the displayed results were evaluated on a sub-sample of manually segmented subjects (12 for WH1 and 12 for WH2) balanced in terms of WMH load and using leave-one-out cross-validation whenever appropriate.*

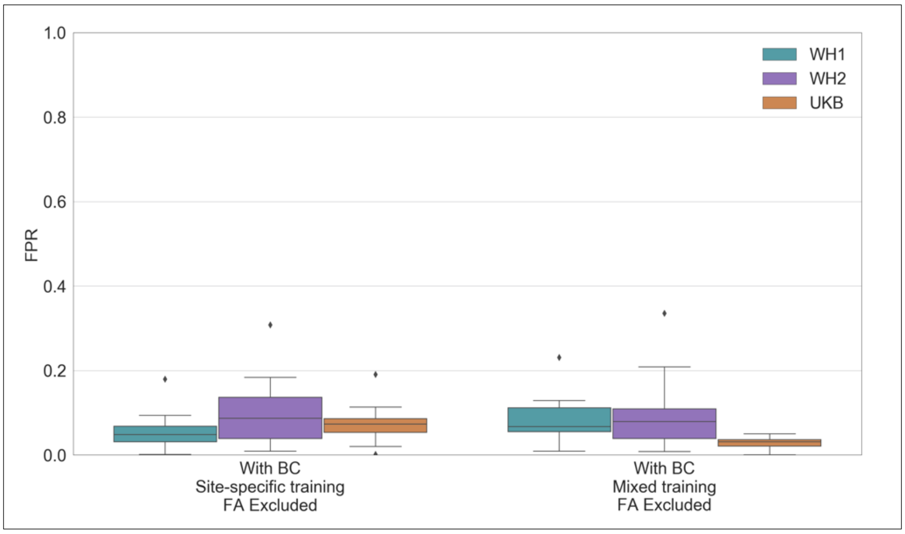


***Figure S5.*** *BIANCA performance – retrospective data merging scenario. Box-plot of the voxel-level False Positive Ratio (FPR) between BIANCA output and the corresponding manual mask for the different analysis options tested during our study (specified on the x axis) All the displayed results were evaluated on a sub-sample of manually segmented subjects (12 for WH1, 12 for WH2 and 12 for UKB) balanced in terms of WMH load and using leave-one-out cross-validation whenever appropriate.*

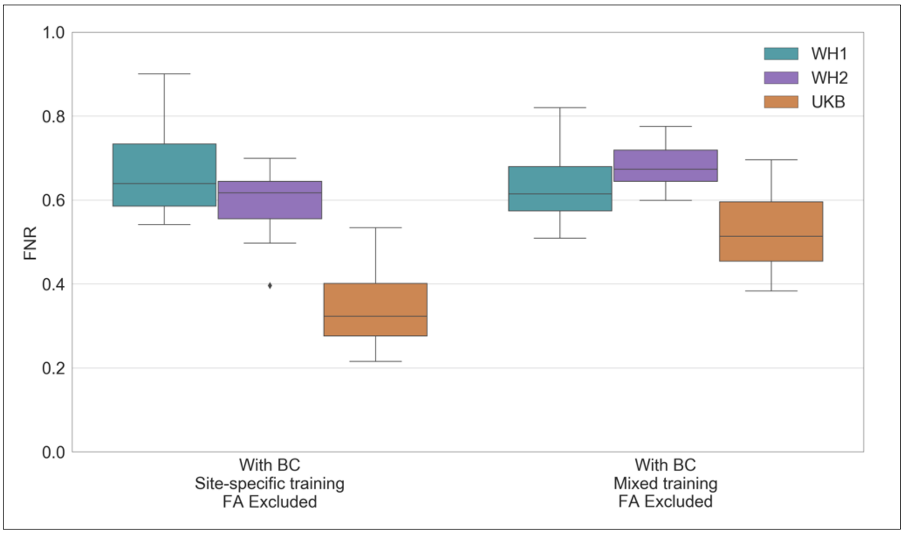


***Figure S6.*** *BIANCA performance – retrospective data merging scenario. Box-plot of the voxel-level False Negative Ratio (FNR) between BIANCA output and the corresponding manual mask for the different analysis options tested during our study (specified on the x axis) All the displayed results were evaluated on a sub-sample of manually segmented subjects (12 for WH1, 12 for WH2 and 12 for UKB) balanced in terms of WMH load and using leave-one-out cross-validation whenever appropriate.*

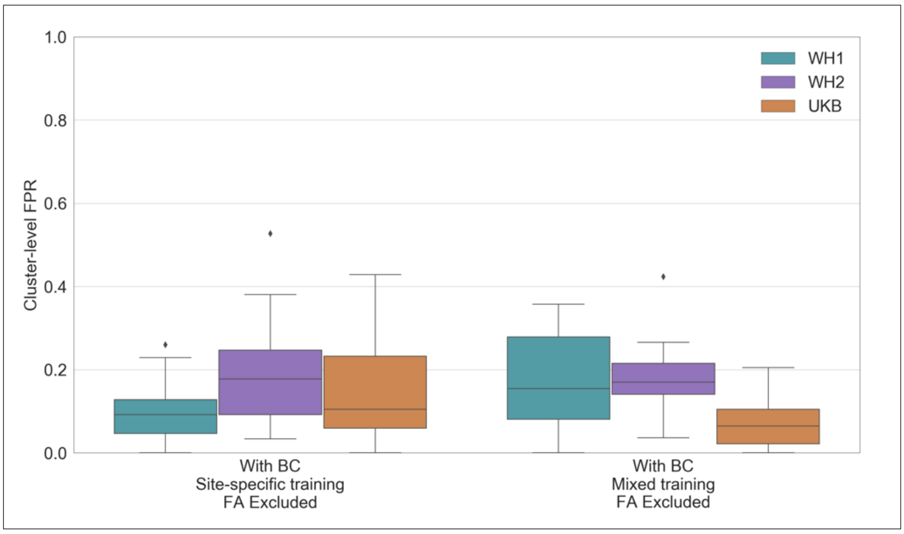

***Figure S7.*** *BIANCA performance – retrospective data merging scenario. Box-plot of the cluster-level FPR between BIANCA output and the corresponding manual mask for the different analysis options tested during our study (specified on the x axis) All the displayed results were evaluated on a sub-sample of manually segmented subjects (12 for WH1, 12 for WH2 and 12 for UKB) balanced in terms of WMH load and using leave-one-out cross-validation whenever appropriate.*

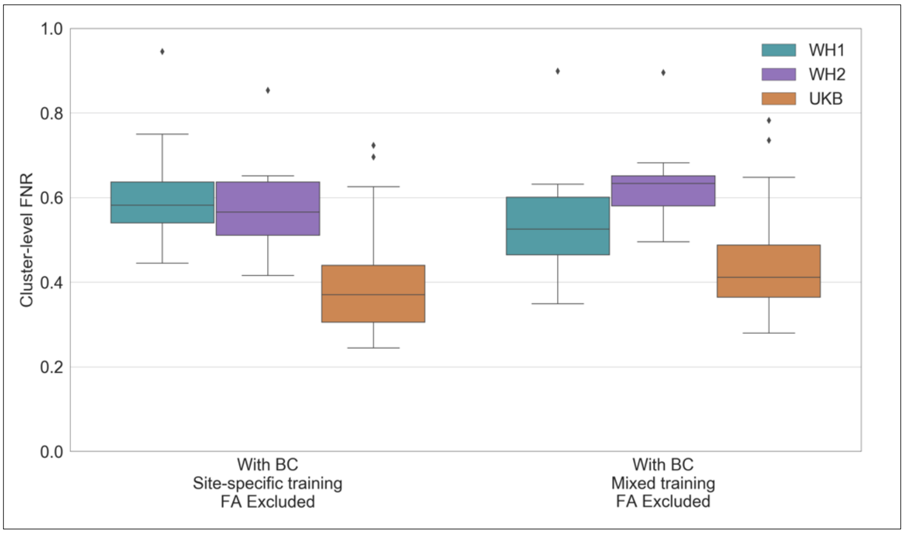


***Figure S8.*** *BIANCA performance – retrospective data merging scenario. Box-plot of the cluster-level FNR between BIANCA output and the corresponding manual mask for the different analysis options tested during our study (specified on the x axis). All the displayed results were evaluated on a sub-sample of manually segmented subjects (12 for WH1, 12 for WH2 and 12 for UKB) balanced in terms of WMH load and using leave-one-out cross-validation whenever appropriate.*

**Table S1.** Effect of rater – Summary of Dice Similarity Index measures calculated for the agreement between manual masks annotated by the raters and BIANCA outputs generated with masks from those raters. Statistical tests performed on data to assess the impact of between- and within-rater variability on the segmentation performance.

| | | R1 vs R2a (between-rater variability) | | R1 vs R2b (between-rater variability) | | R2a vs R2b (within-rater variability) | |
|---|---|---|---|---|---|---|---|
| | | Manual (M1 vs M2a) | BIANCA (B1 vs B2a) | Manual (M1 vs M2b) | BIANCA (B1 vs B2b) | Manual (M2a vs M2b) | BIANCA (B2a vs B2b) |
| Overlap measures | Mean ± std | 0.54 ± 0.18 | 0.43 ± 0.13 | 0.60 ± 0.13 | 0.69 ± 0.10 | 0.54 ± 0.22 | 0.63 ± 0.11 |
| Within-subject analysis: paired t-test | Manual vs BIANCA | < 0.001 *** | | < 0.001 *** | | < 0.001 *** | |

*Results relative to the statistical tests are all reported in terms of p-values: * (< 0.05), ** (< 0.01), *** (< 0.001). Legend: R1 = rater 1, R2a = Rater 2, first rating, R2b = rater 2, second rating (1 year apart from the first rating, blind to first rating), M=manual, B=BIANCA.*

**Table S2.** Elastic Net Regression performance – scanner upgrade and retrospective scenario – Summary of the results in terms of variance explained by the model and by a subset of the features which constituted it. All variables are here reported for the different analysis options tested in our study.

| | | Scanner upgrade scenario | | | | | Retrospective scenario | |
|---|---|---|---|---|---|---|---|---|
| | | Analysis option A | Analysis option B | Analysis option C | Analysis option D | Analysis option E | Analysis option A | Analysis option B |
| **Variance explained by the model** | | 0.243 | 0.161 | 0.207 | 0.173 | 0.125 | 0.244 | 0.133 |
| **Variance explained by the features** | Age | 0.060 | 0.044 | 0.049 | 0.054 | 0.034 | 0.052 | 0.064 |
|  | Sex | 0.014 | 0.014 | 0.004 | 0.014 | 0.006 | ------- | 0.009 |
|  | Width | ------- | ------- | ------- | ------- | 0.000 | ------- | ------- |
|  | Height | 0.012 | 0.006 | ------- | 0.010 | 0.006 | 0.005 | 0.005 |
|  | Body Mass Index | ------- | 0.003 | 0.005 | 0.002 | 0.005 | ------- | ------- |

| | | | | | | | |
|---|---|---|---|---|---|---|---|
| Systolic Blood Pressure | 0.004 | 0.013 | 0.007 | 0.013 | 0.006 | ------- | 0.009 |
| Diastolic Blood Pressure | 0.010 | 0.005 | 0.014 | 0.010 | 0.000 | 0.015 | 0.010 |
| Pulse Rate | ------- | ------- | ------- | ------- | ------- | ------- | ------- |
| Moderate Physical Activity | ------- | ------- | ------- | ------- | ------- | ------- | ------- |
| Vigorous Physical Activity | ------- | 0.001 | 0.002 | ------- | ------- | ------- | ------- |
| CHAM Walk | 0.002 | ------- | ------- | ------- | ------- | 0.006 | 0.003 |
| TV | 0.005 | ------- | ------- | ------- | ------- | ------- | ------- |
| Total Walking Activity | ------- | ------- | ------- | 0.004 | ------- | ------- | ------- |
| Sleep Duration | 0.006 | ------- | 0.002 | ------- | 0.000 | 0.007 | ------- |
| Health Classes | ------- | ------- | ------- | ------- | ------- | ------- | ------- |
| Smoker Status | ------- | ------- | ------- | ------- | ------- | ------- | ------- |
| Smoking Units | ------- | ------- | ------- | ------- | ------- | ------- | ------- |
| Alcohol Status | ------- | ------- | ------- | ------- | ------- | 0.007 | 0.006 |
| Alcohol Units | ------- | ------- | 0.004 | 0.004 | ------- | ------- | ------- |
| Total Medications | ------- | 0.005 | ------- | ------- | ------- | 0.008 | 0.015 |
| Cardiovascular Disease - Medications | 0.013 | 0.008 | 0.011 | 0.019 | 0.009 | ------- | 0.003 |
| Depression - Medications | ------- | ------- | 0.008 | ------- | ------- | ------- | ------- |
| Blood Pressure - Medications | 0.011 | 0.013 | 0.009 | 0.009 | 0.007 | 0.018 | 0.006 |

| | | | | | | | |
|---|---|---|---|---|---|---|---|
| Diabetes | 0.004 | 0.010 | ------- | 0.008 | 0.008 | 0.005 | 0.006 |
| Cardiovascular Disease | ------- | ------- | ------- | ------- | 0.001 | 0.010 | 0.011 |
| Depression | ------- | ------- | ------- | 0.004 | ------- | ------- | ------- |
| Education | ------- | ------- | ------- | ------- | ------- | ------- | ------- |
| Hand – Class | ------- | ------- | 0.003 | ------- | ------- | 0.006 | 0.009 |
| Trail Making Test - A | ------- | ------- | ------- | ------- | ------- | 0.009 | 0.010 |
| Trail Making Test - B | 0.003 | ------- | ------- | ------- | 0.007 | 0.014 | 0.012 |
| Digit Span Backward | ------- | 0.001 | ------- | ------- | ------- | 0.014 | ------- |
| Digit CODing | 0.009 | 0.016 | 0.013 | 0.017 | 0.010 | ------- | ------- |
| Reaction Time | 0.022 | 0.010 | 0.006 | 0.012 | 0.011 | ------- | ------- |
| Scanner | 0.047 | 0.013 | 0.067 | 0.009 | ------- | 0.117 | ------- |

*Options tested in our study are: (I) for the scanner upgrade scenario: (A) without BC, single-site training, FA included; (B) with BC, single-site training, FA included; (C) with BC, site-specific training, FA included; (D) with BC, mixed training, FA included; (E) with BC, mixed training, FA excluded; (II) for the retrospective scenario: (A) with BC, site-specific training, FA excluded; (B) with BC, mixed training, FA excluded. The amount of WMH variance explained by the model is calculated using the R-squared coefficient. The amount of WMH variance explained by the features is reported in the lowest part of the table for all variables.*
